# A refined low-dose murine model of *Mycobacterium ulcerans* infection to assess integrated immune networks in Buruli ulcer pathogenesis

**DOI:** 10.1101/2025.04.23.650345

**Authors:** Stephen Muhi, Isabelle J.H. Foo, Lukasz Kedzierski, Jessica L. Porter, Hayley A. McQuilten, Brian Howden, Katherine Kedzierska, Andrew H. Buultjens, Brendon Y. Chua, Timothy P. Stinear

## Abstract

*Mycobacterium ulcerans*, the causative agent of Buruli ulcer, is a slow-growing zoonotic pathogen with distinctive pathogenesis linked primarily to its toxin mycolactone. Recent research has shown that the *M. ulcerans* infectious dose is very low (<10 colony forming units [CFU]). Buruli ulcer animal infection models traditionally use bacterial challenge doses in the range 10^4^ – 10^6^ CFU; a range orders of magnitude higher than natural infection. These large doses represent an unrealistic challenge for vaccine trials and studies of immunity. Here, we address this issue and describe a murine tail infection model in two genetically distinct mouse strains (BALB/c and C57BL/6) using quality-controlled, *M. ulcerans* challenge doses (10 – 20 CFU and 100 CFU). Over 24-weeks, we assessed host responses to infection by measuring >70 clinical, immunological and microbiological parameters. Principal findings included a 100% infection rate even at the lowest bacterial challenge, but with a dose-dependent delay in lesion onset and disease progression for both mouse strains. Bacterial growth kinetics were similar between mouse strains. There was a difference in immune profiles between mouse strains and between ‘low’ (10 CFU) versus ‘high’ (100 CFU) bacterial challenge doses. C57BL/6 mice exhibited more robust systemic cellular responses and more rapid lesion onset compared to BALB/c mice. There were dose-dependent cytokine and chemokine differences in C57BL/6 mice, while BALB/c mice displayed similar responses across both doses. Antibody responses were only detected late in the infection and were associated with the high-dose inoculum in both strains. Machine learning and other statistical analyses highlighted the importance of activated CD8^+^ T cells and dendritic cells in the immune response to low-dose infection in C57BL/6 mice. Murine low-dose *M. ulcerans* infection models provide confidence for future human Buruli ulcer challenge trials and will inform the development of effective vaccines and therapeutics.

## Introduction

*Mycobacterium ulcerans* is a slow-growing environmental bacterium that causes the neglected tropical skin disease Buruli ulcer (BU) [1]. The distinctive pathology of BU is driven by the *M. ulcerans* toxin mycolactone, a polyketide-derived small molecule with potent immunosuppressive and cytotoxic properties [2]. Observations by noted Australian microbiologist Frank Fenner recorded that 5 – 10 *M. ulcerans* bacilli inoculated into a mouse footpad were sufficient to cause infection, with a long incubation period, up to 5 months [3]. Contemporary research supports these early observations, with an estimated mean incubation period in humans of 4.5 months and an ID_50_ (dose to infect 50%) of just 2 – 3 CFU in BALB/c mice [4–6]. This low infectious dose is like other pathogenic mycobacteria, including *M. leprae* (the causative agent of leprosy) [7] and *M. tuberculosis* [8]. However, *M. ulcerans* vaccine experiments in mice have typically employed quite high bacterial challenge doses, in the range of 10^4^ – 10^6^ colony forming units (CFU) [9]. In Australia, mosquitoes are the vector of *M. ulcerans*, with a relatively low burden of the pathogen associated with the insects, as established by qPCR (294 genomic equivalents [range, 11 – 4,200]) [10]. Thus, findings from prior vaccine- and immunological-related investigations in mice may have confounded protective potential of various experimental vaccines, by overwhelming the host and inducing potentially irrelevant immune responses. Here, we sought to define in depth host integrated immune responses after exposure to low *M. ulcerans* doses across 66 immune parameters, four different time points and two mouse strains.

A major impediment to using low *M. ulcerans* challenge doses has been the technical issue of producing a cell bank with a reliable and reproducible concentration of viable bacteria. *M. ulcerans* is notoriously difficult to handle in the laboratory, as the cells are very hydrophobic and tend to clump [11,12], making preparation of homogeneous cell suspensions difficult. In a previous study, we addressed this bottleneck and described a process to produce a quality-controlled cell bank of low-dose *M. ulcerans* JKD8049 [13], for use in future controlled human infection models [14]. We showed that this inoculum successfully established infection in 20 BALB/c mice, with a 100% attack rate achieved using a dose of only 20 CFU [15]. The overarching goal of the study presented here was to define this quality-controlled cell bank of *M. ulcerans* JKD8049 by dissecting host responses to inform subsequent human challenge trials. The specific aims of our study were to understand the clinical and immunological responses to (i) a narrow range of low *M. ulcerans* challenges doses in (ii) genetically diverse mouse strains (BALB/c and C57BL/6 mice) that are known to have distinct immunological responses to intracellular infection [16]. Pathogen responses are particularly important to examine in BU, as *M. ulcerans* exhibits both intracellular and extracellular phases of infection [17]. Our study was also designed to inform initial dose-finding trials in humans, as the ‘attack rate’ using a very low inoculum (≤ 20 CFU) was hypothesised to be lower in mice with a Th1-biased response to infection (i.e., C57BL/6 mice), particularly in controlling early (intramacrophage) infection, when mycolactone concentration is low [18].

## Materials and methods

### Inoculum preparation

*M. ulcerans* JKD8049, first isolated from a middle-aged male in Point Lonsdale, Victoria, Australia, was prepared and characterised as described previously [13]. In brief, *M. ulcerans* JKD8049 was cultured in Sauton medium (SM) prepared as described previously [13], without the addition of surfactant (e.g., polysorbate/Tween) in order to minimise chemical modification of the mycobacteria. Animal-free and non-genetically modified vegetable peptone broth (Veggietones VG0101, Oxoid, Basingstoke, UK), 10% (v/v), was added as a growth supplement (‘SMVT’). A mechanically filtered suspension of this culture was then cryopreserved at -80°C in 20% (v/v) glycerol. After thawing on ice, the vials were briefly vortexed prior to use. To confirm the final dose injected into mice, a 20 μL volume of a 10-fold dilution series of the suspension was spotted onto Middlebrook 7H10 agar (BD, Sparks, MD, USA) enriched with 10% oleic acid, albumin, dextrose, and catalase (OADC; HiMedia, Mumbai, India). The sample was then drawn into a 1 mL low dead-space (LDS) syringe (B-Braun Omnifix-F, Melsungen, Germany); the sample volume was determined based on the CFU count from other vials produced in the same batch, aiming for two doses: 10 – 20 CFU/mouse and 100 CFU/mouse, in a maximum final volume of 50 μL/dose. A separate 1 mL syringe, containing phosphate buffered saline (PBS) as diluent, was connected to the syringe containing the sample using a luer lock connector (B-Braun Fluid Dispensing Connector, Melsungen, Germany). Mice receiving a ‘high dose’ of *M. ulcerans* JKD8049 were administered mean of 99 CFU in 50 μL for each mouse challenge dose (95% confidence interval [CI] 77 – 124 CFU); mice receiving a ‘low dose’ each received mean of 11.2 CFU (95% CI 9.5 – 12.5 CFU) in 50 μL.

### Animals and ethics statement

BALB/c and C57BL/6 mice were obtained from ARC (Canning Vale, Australia) and housed at the Peter Doherty Institute Bioresources Facilities. All animal work was conducted in accordance with the Australian National Health and Medical Research Council (NHMRC) Code of Practice for the Care and Use of Animals and approved by the Animal Ethics Experimentation Committee (AEC 2023-23710-36088-4) at the University of Melbourne. All mice were monitored twice weekly until a lesion was observed, followed by daily inspection thereafter. All animals were housed in individually ventilated cages, with food and water provided *ad libitum*.

### Mice challenge procedure

Eight-week-old female BALB/c and C57BL/6 mice were infected with *M. ulcerans* JKD8049, with 5 mice of each strain challenged with ‘high dose’ or ‘low dose’ *M. ulcerans.* All mice were lightly anaesthetised with isoflurane and infected subcutaneously in the dorsal surface of the proximal tail, with the needle advanced approximately 3 mm proximally below the skin directly between the lateral tail veins in a 50 μL volume, as previously described [15]. Five control mice of each strain were injected with an identical volume of media only; SMVT prepared as described previously [13] in 20% glycerol (v/v).

### Harvesting of tissues and collection of blood

Spleen, inguinal draining lymph nodes (DLN), tail sections and blood were collected at predefined time points after challenge, with 5 mice of each strain culled at each timepoint (4, 6, 10, 14 and 28 weeks after challenge, or at the onset of ulceration). Spleens and DLNs were collected in 3 mL Roswell Park Memorial Institute (RPMI; Life Technologies) medium containing 2% foetal calf serum (FCS) (v/v) and passed through 70 µm cell sieves (Miltenyi Biotech) to obtain single cell suspensions. Splenic-derived cell suspensions were further treated with 0.15 M NH_4_Cl and 17 mM Tris-HCl at pH 7.2 for 5 minutes at 37°C to lyse red blood cells and washed with medium prior to analysis. Blood was collected from mice immediately post-mortem from a terminal cardiac bleed. Blood was then placed on ice for 4 – 6 hours, after which clots were removed manually. The samples were then centrifuged at 2,000 rpm for 5 minutes at 4°C, and supernatant collected for cytokine and chemokine analysis, and antibody responses. Tail tissue was prepared for analysis as described previously [4]. Briefly, tail sections without lesions were harvested by dissecting the proximal third of the tail and ulcerated tails were dissected with a small margin on either side of the lesion. For homogenisation, tail sections were placed in 600 – 1000 µl of PBS and processed by bead beating using ∼0.5 g 2 mm glass beads in a high-speed tissue-disruptor at 6,500 rpm (Precellys 24, Bertin Technologies, France). Homogenised tissue samples were stored at -80°C.

### Immunophenotypic staining

Cells were stained with combinations of fluorochrome-conjugated antibodies (clone; catalogue number; dilution). BD Biosciences: anti-CD8α-PerCyP Cy5.5 (53-67; 551162; 1:200), anti-CD4-BV650 (RM4-5; 563747; 1:200), anti-CD25-PECF594 (PC61; 562694; 1:200), anti-CD45-PerCP (30-F11; 557235; 1:200), anti-CD38-BV711 (90; 740697; 1:200), anti-CD138-PE (281-2; 553714; 1:400), anti-CD11b-BV605 (M1/70; 563015; 1:600), anti-SigLecF-PECF594 (E50-2440; 562757; 1:1200), anti-Ly6G-PECy7 (1A8; 560601; 1:1200), anti-CD19-BV605 (1D3, 563148, 1:200), anti-CD27-PE (LG.3A10, 558754, 1:200), anti-CD49b-APC (DX5, 560628, 1:100), anti-I-A/I-E-AF488 (M5/114, 562352, 1:200), anti-iNOS-FITC (6/iNOS/NOS Type II, 610331, 1:100), anti-KLRG1-APC (2F1, 561620, 1:200), anti-NK1.1-AF700 (PK136, 560515, 1:100), and anti-TCRγδ-BV421 (GL3, 562892, 1:200).

BioLegend: anti-CD62L-BV570 (MEL-14; 104433; 1:200), anti-CD279-BV785 (29F.1A12; 135225; 1:200), anti-CD69-PECy7 (H1.2F3; 104512; 1:200), anti-GL7-PerCyPCy5.5 (GL7; 144610; 1:200), anti-IgD-APC Cy7 (11-26c.2a; 405716; 1:200), anti-CD3-BV785 (145-2C11; 100355; 1:200), anti-F4/80-APC (BM8; 123116; 1:150), anti-CD45R-BV650 (RA3-6B2, 103241, 1:200), anti-CD107a-BV421 (1D4B, 121617, 1:100), anti-CD11c-BV650 (N418, 117339, 1:200), anti-CD44-APC Cy7 (IM7, 103028, 1:200), anti-CD64-BV421 (X54-5/7.1, 139309, 1:100), and anti-Ly6C-BV570 (HK1.4, 128030, 1:200).

eBiosciences: anti-I-A/I-E-APC-eFlour780 (M5/114.15.2; 47-5321-82; 1:200), anti-Arg1-AF700 (A1exF5, 56-3697-82, 1:100), anti-CD206-PE (MR6F3, 12-2061-82, 1:100), and anti-CD3-AF700 (eBio500A2, 56-0033-82, 1:200).

Cell viability was determined by staining with Live/Dead-Aqua 525 (L34966A, ThermoFisher, 1:800). Antibody staining was performed at 4°C in the dark. Cells were fixed with 1% paraformaldehyde before analysis by flow cytometry. Samples were subsequently acquired on a BD LSR Fortessa (BD Biosciences) flow cytometer and data analysed by FlowJo Software (Tree Star Inc., USA). Gating strategies for flow cytometry data are shown in Fig S1.

### Establishing colony counts

A 300 μL volume of tail tissue homogenate was decontaminated with 300 μL of 2% NaOH (v/v) and incubated at room temperature for 15 minutes. The preparation was then neutralised with 50 μL of 10% orthophosphoric acid (v/v). 300 μL of this mixture was then diluted in 300 μL PBS and CFUs were determined by spot plating as described previously [4]. In brief, 6 x 5 μl volumes of serial 10-fold dilutions (10^−1^ to 10^−5^) of tissue preparation were spotted onto 7H10/OADC agar plates with a 5 x 5 grid. A second plate (i.e., technical replicate) was also spotted in case of contamination. The spots were allowed to dry, loosely wrapped in a plastic bag and incubated for 12 weeks before final colony count. Plates were checked every two weeks.

### Histopathology

In groups of mice without any visible lesion, 2 of 5 mice tails were dissected for histology using the entire proximal third of the tail. In mice with a visible lesion, the segment of involved tail tissue was divided into two equal sections. Samples of excised tissue were fixed in neutral buffered formalin, decalcified in 10% formic acid for 48 hours and embedded in paraffin for histopathological examination. Tissue sections were then stained with hematoxylin and eosin (HE) and Ziehl-Neelsen (ZN) for light microscopy. Digital images were captured using a Pannoramic SCAN II (3D Histech), using a Plan-Apochromat 20×/NA 0.8 objective (Carl Zeiss Microscopy) and a Grasshopper 3 CCD monochrome camera (Point Grey).

### DNA extraction and PCR

DNA was extracted from homogenised tail tissue using a DNeasy PowerSoil kit (Qiagen). Control mice were included to monitor potential PCR contamination in addition to no-template negative PCR controls. IS*2404* quantitative PCR (qPCR) was performed using technical duplicates as described previously [19]. IS*2404* cycle threshold values were converted to genomic equivalents as described previously [4]. In brief, cycle thresholds were converted using a reference standard curve calculated using dilutions of genomic DNA from *M. ulcerans* JKD8049 [4].

### Serum cytokines and chemokines analysis

Cytokine and chemokine levels in mice sera were analysed using the Mouse LEGENDplex™ MU Cytokine Release Syndrome Panel (13-plex) kit (Biolegend) according to manufacturers’ instructions.

### Antibody responses

*M. ulcerans* whole cell lysate (WCL) was prepared using *M. ulcerans* JKD8049 cultured on solid agar for > 12 weeks. Culture material was scraped into 2 mL vials containing ∼ 0.5 g 2 mm glass beads and 1 mL of sterile PBS and homogenised in a high-speed tissue-disruptor. Homogenised lysate was then centrifuged at 4,000 rpm for 5 minutes and the supernatant was transferred into a separate container. The final protein concentration (0.4 mg/mL) was determined using a Qubit Protein Assay (Invitrogen), as per manufacturer’s instructions for use.

Enzyme-linked immunosorbent assays (ELISAs) were performed on Nunc Immuno MaxiSorp PS flat-bottom 96 well plates (Thermo Scientific). Each plate well was coated with 50 µL of PBS containing 5 µg/ml of WCL and incubated overnight at 4°C. The following day, WCL was discarded and 100 µl of PBS containing w/v 10% bovine serum albumin (BSA_10_PBS) was added to each well. This was incubated for 1 hour at room temperature (RT). The BSA_10_PBS was then discarded, and the plate washed 4 times with PBS containing v/v 0.05% Tween (PBST) and once with PBS. Plasma was added to the wells in a log_0.5_ dilution series in PBS containing v/v 0.05% Tween and w/v 5% PBS (BSA_5_PBST), and incubated for 2 hours at RT. Following discard of well contents, each plate was washed as described. For immunoglobulin detection, 50 µl of polyclonal rabbit anti-mouse Ig-HRP (Dako) was added to each well at a 1:2000 dilution in BSA_5_PBST and incubated for 1 hour at RT. The content of each plate was then discarded and washed 7 times with PBST and once with PBS followed by the addition of 100 µl of tetramethylbenzidine (TMB; Sigma) and allowed to develop for 10 minutes before 100 µl of 1 M orthophosphoric acid was added. Plates were then read immediately at an absorbance wavelength of 450 nm on a Multiskan MS multiplate reader (Labsystems). Endpoint titers were determined by interpolation from a sigmoidal curve fit (all R-squared values >0.95; GraphPad Prism 10) and expressed as the reciprocal of the highest dilution of plasma required to produce an absorbance value of 0.8. Intra-experimental measurements were normalised using a positive control plasma sample included on each plate.

### Statistical analysis

Statistical analysis was performed using GraphPad PRISM (version 10.1.1). Continuous variables are reported as mean/standard deviation or 95% confidence interval, as relevant. Comparisons between groups were performed using Student’s t-test (for difference between means) and illustrated using Kaplan-Meier curves using the log-rank test, with statistical significance if p < 0.05. Univariate Mann-Whitney U tests were performed in python using SciPy (version 1.9.1) to compare individual immune features between the low and high dose groups. To adjust for false discovery due to multiple comparisons, from each group of highly correlated variables (defined as correlation coefficient > 0.8), only one representative variable was retained in order to minimise redundancy. The Benjamini– Hochberg procedure was then applied, with statistical significance defined as p < 0.05.

Locally Weighted Scatterplot Smoothing (LOWESS) regression was conducted in Python (v3.10.6) using pandas (v1.4.2) and numpy (v1.23.3), with visualisations created in matplotlib (v3.7.1). Linear models of log-transformed variables were constructed in R (v4.2.1) and plotted with ggplot2 (v3.3.5). Heatmaps were generated using seaborn (v0.12.2), where values for each immune parameter were normalised across all timepoints and groups (control, low and high dose challenge).

Random Forest classifiers were applied to evaluate the ability of immune parameters to distinguish between low and high dose challenge groups using sklearn (v1.1.2). The data were divided into 10 stratified folds and feature importances were averaged across folds. Classifier performance was assessed by calculating the area under the receiver operating characteristic curve (AUC), with results aggregated across all folds. To establish a null distribution, the analysis was repeated 100 times with randomised class labels and AUC scores from these runs were compared to the actual data using density plots.

Pearson correlation coefficients were calculated in R version 4.4.1 using the psych (v2.4.6.26) package function corr.test, with p-values representing the Benjamini-Hochberg false discovery rate adjustment for multiple t-tests. Pairwise correlation plots were made using ggplot2 version 3.5. Correlation matrices were made using corrplot (v0.95).

## Results

### Low challenge doses of M. ulcerans JKD8049 lead to tail lesion formation and progression of clinical disease in both C57BL/6 and BALB/c mice

To compare the pathology and immunology of different *M. ulcerans* dosing strategies, we challenged C57BL/6 and BALB/c mice with a low and high dose of *M. ulcerans*, corresponding to an average 11.2 CFU (95% CI 9.5 – 12.5 CFU) and 99 CFU (95% CI 77 – 124 CFU) (Fig S2A), respectively.

We then followed responses to infection at 4, 6, 10 weeks following challenge, and at the onset of tail ulceration. Negative control mice were challenged with culture media only. In both C57BL/6 and BALB/c mice receiving either dose, typical BU lesions were observed in all mice; namely, lesions began as a small patch of erythema with hair loss that with progressively enlarging induration and eventual ulceration (Fig 1A).

**Figure 1.**
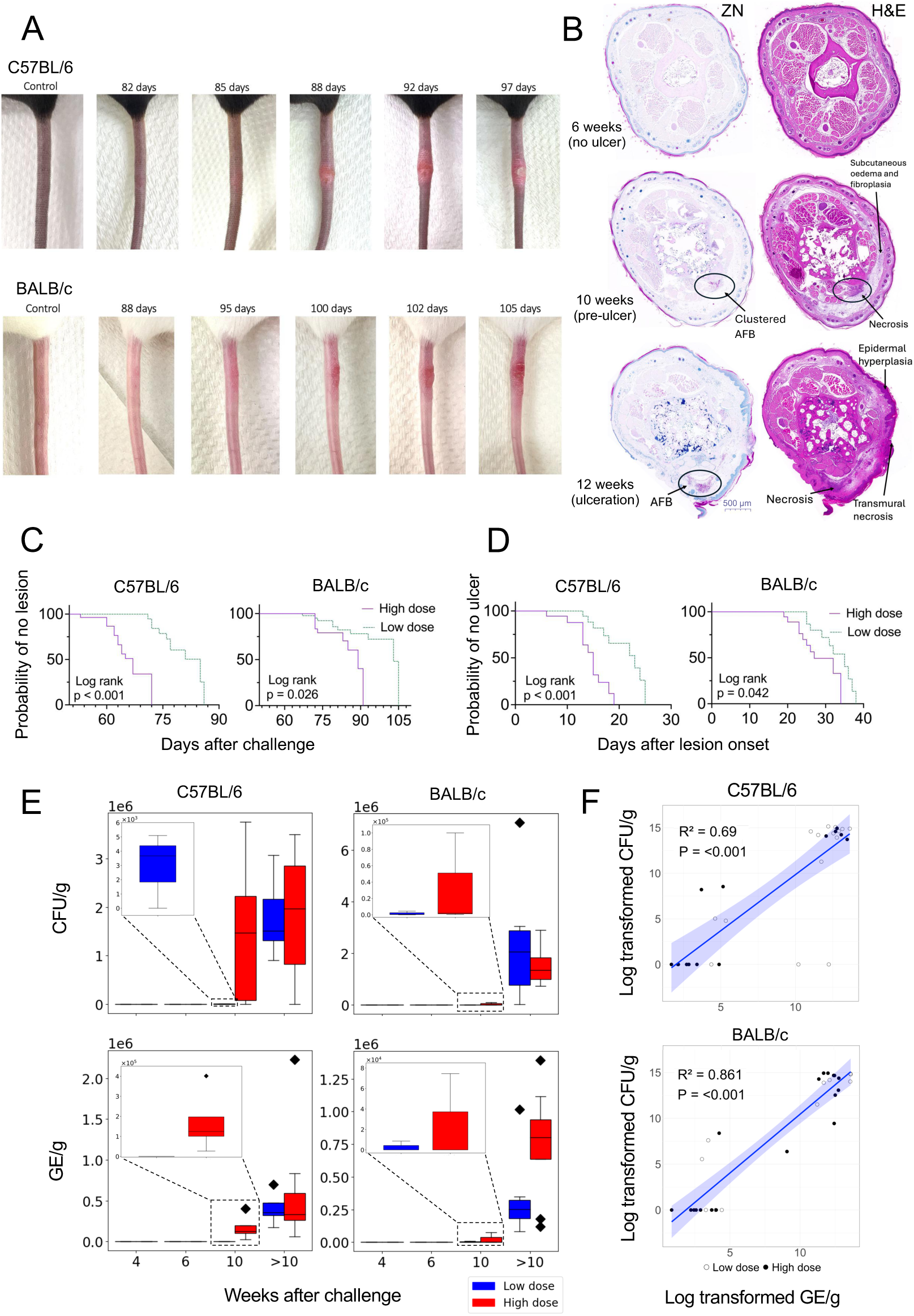
Clinical and microbiological features of low-dose infection over time. **A** Typical progression of tail lesions in female C57BL/6 (top row) and BALB/c (bottom row) mice challenged subcutaneously with ∼ 100 CFU of *M. ulcerans* JKD8049. **B** Histopathological features of C57BL6 mouse tail imaged on day 42, 70 and 84 following challenge with ∼ 100 colony forming units (CFU) of *M. ulcerans* JKD8049. Tail sections were stained with Ziehl-Neelsen (ZN) acid-fast stain and hematoxylin and eosin (H&E) stain; representative images are shown. Bar represents 500 μm length at magnification of 2x. Circled area shows presence of acid-fast bacilli (AFB) and arrows point to areas of histopathology. **C** Differences in the time to lesion onset in BALB/c and C57BL/6 mice depending on the challenge ‘dose’ received (∼10 CFU challenge dose: n=10 per group; ∼100 CFU challenge dose: C57BL6 n=15, BALB/c n=10). **D** Time from lesion onset to ulceration in these mice (∼10 CFU challenge dose: n=10 per group; ∼100 CFU challenge dose: C57BL6 n=12, BALB/c n=10). Comparisons between groups were performed using the log-rank test. **E** Bacterial growth over time in CFU/g and genomic equivalents (GE)/g. **F** Relationship between the log transformed CFU/g and GE/g in both mouse groups (C57BL/6 top, BALB/c bottom); comparisons between groups were performed using linear models constructed in R (v4.2.1) on log-transformed variables (selected using the bestNormalize package). Model fit was evaluated using R², while the significance of association was determined using p-values from t-tests on the model coefficients.

Histopathology of tail sections for C57BL/6 mice at 10 weeks post low dose challenge showed evidence of pre-ulcerative disease, with mild focal subcutaneous oedema and scanty lymphocytic infiltration. No acid-fast bacilli (AFB) were detected by Ziehl–Neelsen (ZN) staining at this time point (Tables S1 and S2). These inflammatory features were not as apparent for similarly challenged BALB/c mice (Tables S3 and S4). In C57BL/6 mice that received the high dose, the pathology in tail sections was prominent at earlier timepoints, and typified by moderate multifocal to diffuse subcutaneous oedema. Two of five of these mice at the 10-week time-point showed mononuclear cell infiltration, while three mice had a mixed inflammatory infiltrate associated with vasculopathy and skeletal myofiber necrosis (Fig 1B). One C57BL/6 mouse at this timepoint also had intracellular AFB visible within macrophage cytoplasm. By comparison, in the low-dose C57BL/6 mice examined 10 weeks after infection, the two mice examined histologically both had mild subcutaneous oedema and scanty lymphocytes, with no AFB visible. At later timepoints (>10 weeks) there was visible ulceration in both dose groups and mouse strains. Common histological features in these mice included subcutaneous oedema with a mixed infiltrate, vasculopathy, myonecrosis, and abundant AFB. Of note, BALB/c mice culled late in the low dose challenge group also showed chronic, active inflammation that was not observed in any mice given a high dose. Moreover, in all mice with visible lesions, 3 of 25 mice in the high dose cohorts had contiguous AFB detected in the underlying bone marrow. This phenomenon was not detected in any of the mice with visible lesions (n=19) challenged with the low dose of *M. ulcerans* (Fig S2B, Tables S1 and S3).

### Low challenge doses of M. ulcerans result in a longer incubation period and slower progression of clinical disease in both BALB/c and C57BL/6 mice

There was a clear dose-response relationship in both mouse strains. Low-dose mice, irrespective of strain, developed lesions later than those receiving the high dose (Fig 1C). Mice receiving a low dose of *M. ulcerans* also had a slower progression of clinical disease (the time from lesion onset to ulceration), regardless of mouse strain (Fig 1D). Additionally, despite infecting with identical bacterial doses, disease onset and disease progression was significantly slower in BALB/c mice relative to C57BL/6 mice (Fig S2C, S2D).

In C57BL/6 mice, the mean time-to-lesion-onset (*i.e.* incubation period) was 64 days for the high dose compared to ∼77 days following a low dose (delay in symptom onset was 13 days) (Table 1). Similarly, visible lesions were also observed in all infected BALB/c mice, with mean time-to-lesion-onset of ∼79 days in those that received a high dose compared to ∼84 days for the low dose (delay in symptom onset ∼5 days).

**Table 1.** Clinical parameters of mice allowed to progress to clinical disease, depending on mouse type and dose of *M. ulcerans* JKD8049.

|  | Dose group and attack rate (%) |  | Mean incubation (days) (95% CI) | Mean time from lesion onset to ulcer (days) (95% CI) | Delay in incubation period (days) | Delay in time from lesion onset to ulceration (days) |
| --- | --- | --- | --- | --- | --- | --- |
| C57BL/6 | High dose | 15/15<br>(100%) | 64.1<br>(53.0 – 72.0) | 13.4<br>(11.7 – 16.1) |  |  |
| C57BL/6 | Low dose | 10/10<br>(100%) | 77.1<br>(71.0 – 86.0) | 19.6<br>(16.2 – 23.0) | 13.0 | 5.7 |
| BALB/c | High dose | 10/10<br>(100%) | 79.1<br>(72.0 – 89.0) | 25.3<br>(21.9 – 28.7) |  |  |
| BALB/c | Low dose | 10/10<br>(100%) | 83.8<br>(74.6 – 93.0) | 31.4<br>(27.8 – 35.0) | 4.7 | 6.1 |

Upon appearance of visible lesions, the mean time-from-lesion-onset progressing to ulceration in C57BL/6 mice was ∼14 days following a high dose and ∼20 days in those that received a low dose (delay in ulceration ∼ 6 days) (Table 1). In comparison, the mean time from lesion onset to ulceration in BALB/c mice was ∼25 days in high dose mice and ∼32 days for low dose (delay in ulceration ∼ 6 days). These results demonstrate dose-dependent disease outcomes with a high dose exhibiting more rapid appearance of lesions and development of ulcers in both mouse strains. There was also a faster rate of symptom progression in C57BL/6 mice compared to BALB/c mice, irrespective of the dose.

Of C57BL/6 mice that developed ulceration, some weight gain was observed compared to uninfected controls but this change was not statistically significant (Table S5). No weight change was observed for BALB/c mice (Table S6). All infected mice appeared otherwise healthy and did not appear to groom or attend to their lesions, and no secondary lesions were observed.

### Exponential M. ulcerans growth occurs in both BALB/c and C57BL/6 mice after both high and low challenge doses

To characterise bacterial growth dynamics during infection, IS*2404* qPCR and colony forming unit (CFU) counts were performed on mouse tail homogenates. *M. ulcerans* was detected by culture in tail tissue from high dose C57BL/6 mice from 6 weeks post inoculation (Table 2, Fig 1E). Despite the absence of any visible lesion, two of three samples collected at 6 weeks resulted in bacterial colony growth (mean 136 CFU/g). In contrast, low dose C57BL/6 mice were culture positive from 10 weeks, at which time two of three samples were positive in the absence of any lesion (mean 4.39 × 10^3^ CFU/g). Tail tissue from BALB/c mice were culture positive from 10 weeks, where all three tails tested from the high dose challenge cohort were positive (mean 3.41 × 10^4^ CFU/g) compared to two of three tails in the low dose challenge group (mean 2.47 × 10^3^ CFU/g). All mice culled >10 weeks after challenge had ulcerative disease and bacterial counts were high in these mice (mean 2 × 10^6^ CFU/g) irrespective of low or high dose inoculum. There were no differences in *M. ulcerans* growth kinetics between BALB/c and C57BL/6 mice (Fig S2E). The lower dose resulted in later time to detection of viable bacilli in both mouse strains. Using *M. ulcerans* qPCR that is more sensitive and less resource-intensive than culture, we observed an exponential pattern of growth by plotting genomic equivalents (GE) per gram of tail tissue over time (Fig 1E, Fig S2F). There was a significant positive correlation between CFU/g and GE/g in both mouse strains (Fig 1F).

**Table 2.** *M. ulcerans* JKD8049 CFU count and genomic equivalents in mouse tail homogenate.

| Weeks after challenge | No. culture positive | No. with lesion | Mean CFU/g | SD | Mean GE/g | SD |
| --- | --- | --- | --- | --- | --- | --- |
| C57BL/6 High dose |  |  |  |  |  |  |
| 4 | 0 of 3 | 0 | N/A | N/A | 83 | 16 |
| 6 | 2 of 3 | 0 | 136 | 23 | 128 | 69 |
| 10 | 4 of 5 | 5 | 1.88E+06 | 1.54E+06 | 1.70E+05 | 1.43E+05 |
| > 10 | 10 of 10 | 10 | 1.64E+06 | 1.15E+06 | 2.96E+05 | 1.75E+05 |
| C57BL/6 Low dose |  |  |  |  |  |  |
| 4 | 0 of 3 | 0 | N/A | N/A | 16 | 12 |
| 6 | 0 of 3 | 0 | N/A | N/A | 11 | 5 |
| 10 | 2 of 3 | 0 | 4.39E+03 | 1.01E+03 | 114 | 68 |
| > 10 | 7 of 10 | 10 | 2.21E+06 | 1.00E+06 | 5.90E+05 | 6.29E+05 |
| BALB/c High dose |  |  |  |  |  |  |
| 4 | 0 of 3 | 0 | N/A | N/A | 48 | 8 |
| 6 | 0 of 3 | 0 | N/A | N/A | 53 | 28 |
| 10 | 3 of 3 | 0 | 3.41E+04 | 5.71E+04 | 2.49E+04 | 4.30E+04 |
| > 10 | 10 of 10 | 10 | 2.31E+06 | 2.23E+06 | 9.24E+05 | 5.17E+05 |
| BALB/c Low dose |  |  |  |  |  |  |
| 4 | 0 of 3 | 0 | N/A | N/A | 19 | 19 |
| 6 | 0 of 3 | 0 | N/A | N/A | 10 | 2 |
| 10 | 2 of 3 | 0 | 2.47E+03 | 2.67E+03 | 2.92E+03 | 4.95E+03 |
| > 10 | 10 of 10 | 10 | 2.20E+06 | 2.03E+06 | 3.08E+05 | 2.63E+05 |

CFU: colony forming units; SD: standard deviation; GE: genomic equivalents.

In C57BL/6 mice, the higher dose correlated with faster time-to-lesion onset (Fig 1C) and more rapid progression of clinical disease (Fig 1D). In BALB/c mice, although a similar pattern emerged, the resolution between high and low challenge doses was less apparent, suggesting that clinical outcomes are less readily distinguished between the two doses in BALB/c mice (Fig 1C, 1D). In C57BL/6 mice, despite mice progressing faster to visible clinical disease following the high dose, there were similar or greater bacteria in mice at the time of ulceration that received the low dose (Fig S3A, S3B). This suggests that pathology at these clinical endpoints may be driven by non-bacterial processes in C57BL/6 mice, a pattern which was not as apparent in BALB/c mice.

### C57BL/6 mice demonstrate an early splenic response to low-dose infection, followed by a late response in draining lymph node tissue

We next investigated immunological responses over time systemically, as captured in splenic tissue, and peripherally in the inguinal draining lymph nodes (DLNs) that are proximal to the site of challenge at the base of the tail. We first analysed immune cell populations in the spleen and DLNs over the course of infection progression. Compared to uninfected C57BL/6 mice, increased immune cell numbers in the spleen were apparent 4 – 6 weeks following low dose challenge, but decreased to baseline levels from 10 weeks onwards, concomitant with a rise in bacterial numbers (Fig 2A). A similar peak in cell numbers was observed in those mice that received the high dose, albeit occurring later at 6 weeks, consistent with an earlier increase in bacterial burden at 10 weeks. In contrast, cell numbers in the DLNs did not peak until 10 weeks after challenge, most prominently in the low-dose group, and like observations in the spleen, DLN cell numbers subsided as bacterial concentrations increased. However, in BALB/c mice challenged with either dose, there were no discernible changes in splenic and DLN cell numbers over the course of the following 14 weeks, despite increases in the bacterial burden at the humane endpoint, with the exception of the DLNs from those mice that received the higher dose, which saw an increase in cell numbers at 10 weeks post-challenge (Fig 2B).

**Figure 2.**
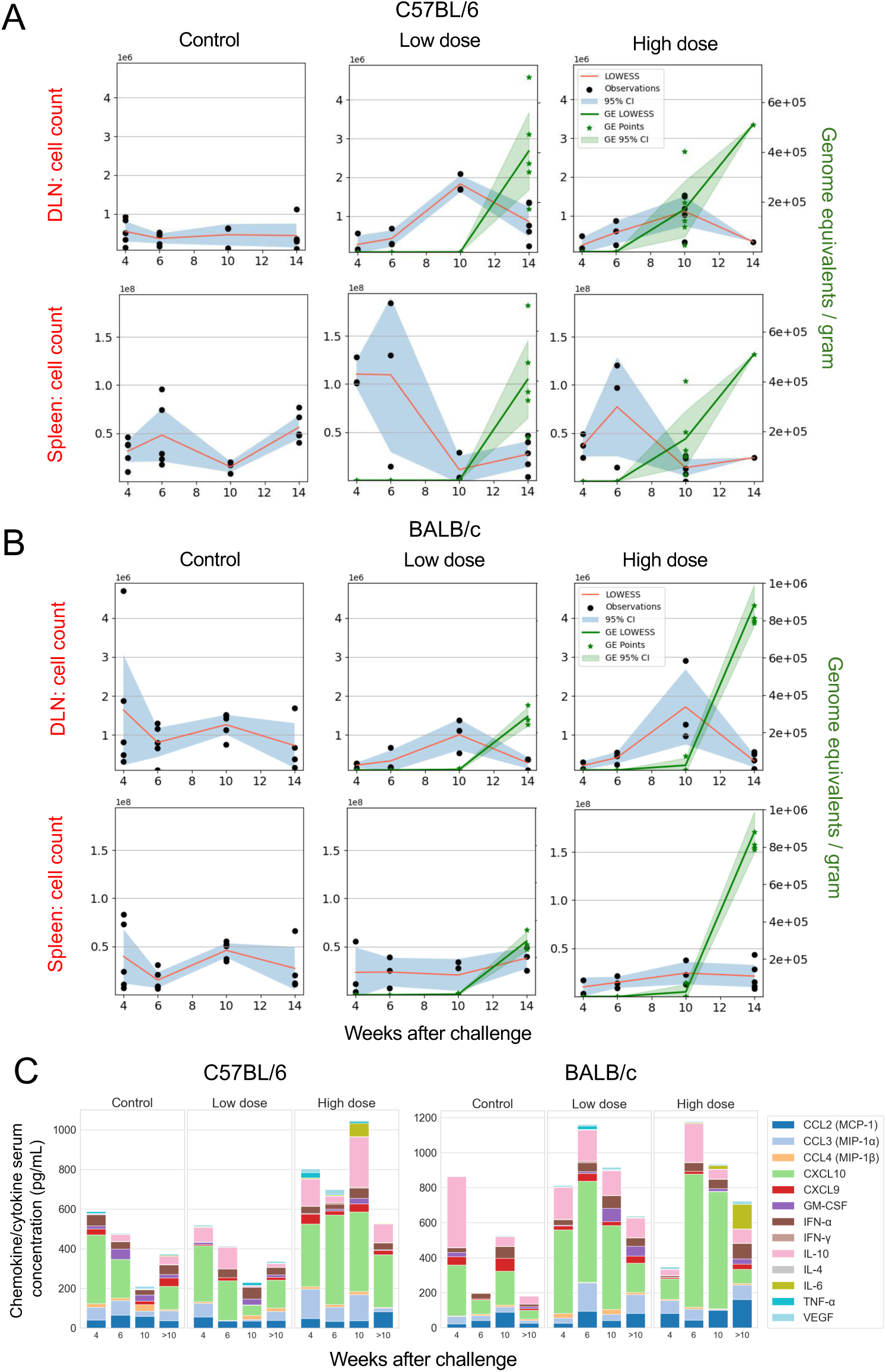
Cellular and cytokine/chemokine features over time. **A** Total cell counts observed in C57BL/6 and **B** BALB/c mice at 4, 6 and 10 weeks after challenge (as per protocol), in splenic tissue and draining lymph nodes following either media-only ‘sham’ (control mice, left column) or infectious challenge. Mice which received a ‘low dose’ challenge are visualised in the centre column, and mice that received a ‘high dose’ of *M. ulcerans* are visualised in the right column. Images also include mice culled 14 weeks after challenge, at which time mice had reached the human end-point (ulceration). The blue colour represents the 95% confidence interval around the Locally Weighted Scatterplot Smoothing (LOWESS) line (in red), and the green colour represents the genomic equivalents (GEs) LOWESS line and 95% confidence interval (C57BL/6 mice controls: week (W) 4 – 6 n=5 per group, W10 n=3, W14 n=5; C57BL/6 ∼10 CFU dose: W4 n=4, W6 – 14 n=5 per group; C57BL/6 mice ∼100 CFU dose: W4 – 10 n=5 per group, W14 n=1; BALB/c mice controls: W4 – 6 n=5 per group, W10 – 14 n=4 per group; BALB/c mice ∼10 CFU dose: W4 – 10 n=5 per group, W14 n=3; BALB/c mice ∼100 CFU dose: W4 n=4, W6 – 14 n=5 per group). **C** Serum cytokine and chemokine concentrations over time across a maximum of 140 days were assayed by LegendPlex; C57BL/6 mice controls: W4 – 6 n=5 per group, W10 n=3, W>10 n=12; C57BL/6 mice, ∼10 CFU dose: W4 n=4, W6 – 10 n=5 per group, W>10 n =10; ∼100 CFU dose: W4 –10 n=5 per group, W>10 n=10; BALB/c mice controls and both dosing groups: W4 – 10 n=5 per group, W>10 n=10.

### Cytokine/chemokine responses are dose-dependent and more prominent in C57BL/6 mice

An analysis of 13 serum cytokines and chemokines in C57BL/6 mice demonstrated that only IL-6 was greater in high dose mice than low dose (p=0.005) (Fig. 2C, Fig S4A, Table S7). In BALB/c mice, the pattern of cytokine/chemokine responses was similar between the doses, peaking 6 weeks following infection, then subsiding (Fig 2C). IFN-γ was significantly greater in the high dose mice compared to the low dose early in the course of infection (p=0.045). IL-10 was significantly greater in the high dose mice compared to low dose and control mice 10 weeks after challenge (p=0.045, p=0.044 respectively) (Fig 2C, Fig S4B, Table S7). Given the role of IL-10 in regulating immune responses in general and the elevated IL-10 responses with inversely low IFN-γ expression in humans with BU, the levels of these cytokines may at least partially explain the reduced inflammatory response observed in BALB/c mice [20].

### Differential systemic versus local innate and adaptive cellular immune responses between mouse strains and between doses

Innate and adaptive immune cell populations over time were further analysed to understand immune features influencing differences between dosing groups. In C57BL/6 mice, there were several adaptive immunity cell populations that were significantly different in number between dosing groups and controls (Fig 3A). In splenic tissue of C57BL/6 mice after low dose challenge at 4 weeks, when compared to control mice, there were increased numbers of CD8^+^ and CD4^+^ T cells, T_reg_ cells, B cells and plasma cells (all p=0.016) (Table S7). In splenic tissue of high dose C57BL/6 mice, these early differences were not apparent, and in some cases, numbers of these adaptive immune cells were significantly less than those observed in low dose mice, particularly CD8^+^ T cells, CD4^+^ T cells, T_reg_ cells, B cells (all p=0.016) and plasma cells (p=0.032) at 4 weeks (Table S7).

**Figure 3.**
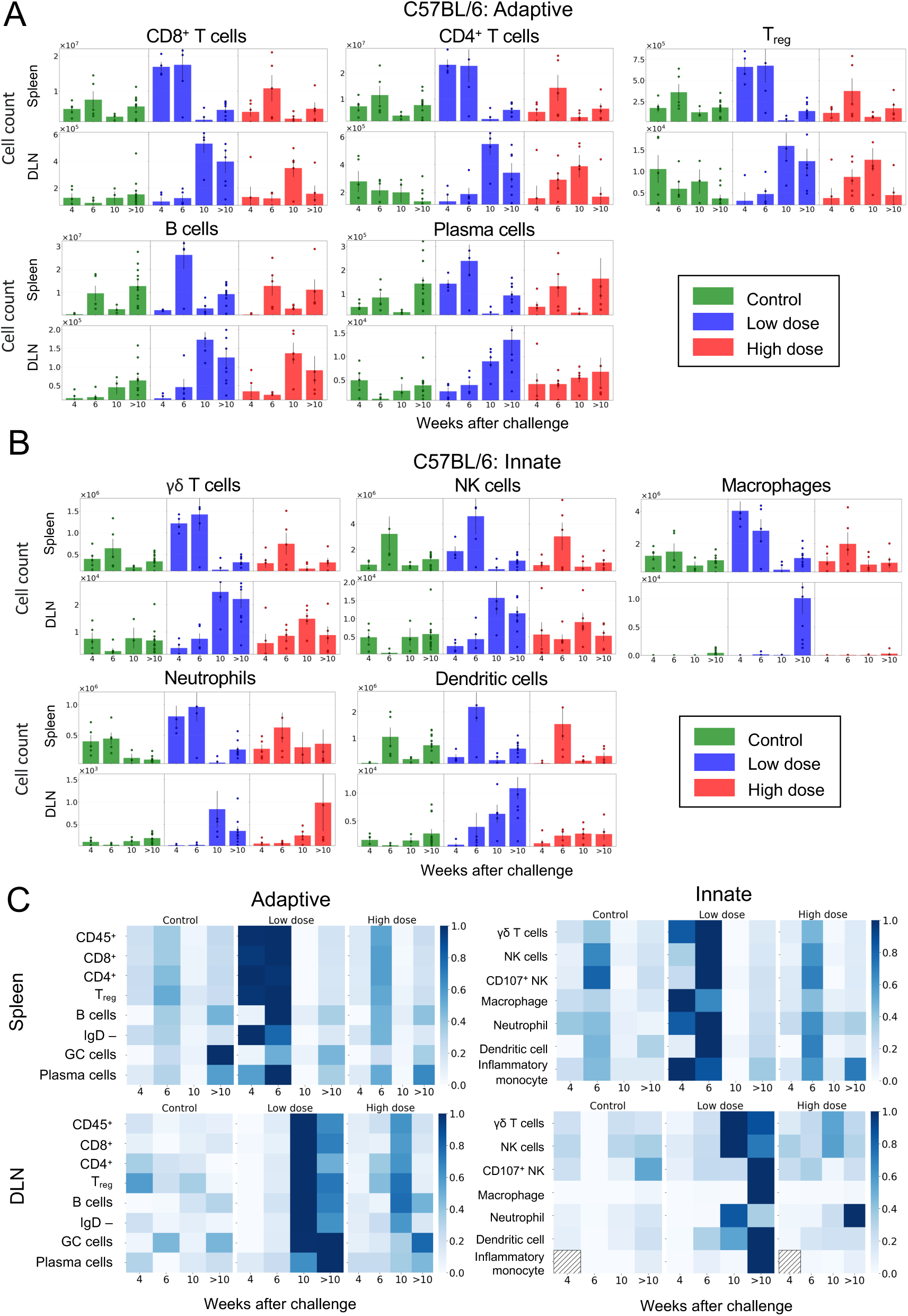
Total cell counts of adaptive and innate immune features over time in C57BL/6 mice following low-dose *M. ulcerans* infection. **A** Adaptive and **B** innate immune features over a maximum of 15 weeks using absolute numbers of effector CD8^+^ T cells (CD8^+^ CD62L^lo^ CD44^hi^), effector CD4^+^ T cells (CD4^+^ CD62L^lo^ CD44^hi^), T_reg_ cells (CD4^+^ CD25^+^ CD62L^+^), B cells (CD19^+^ B220^+^), plasma cells [antibody secreting cells] (IgD-B220^lo^ CD138^+^), γδ T cells (TCRγδ CD3^+^), NK cells (CD49b^+^ NK1.1^+^), macrophages (CD64^+^ F4/80^+^), neutrophils (CD11b^+^ Ly6G^+^) and dendritic cells (CD11c^+^MHCII^+^). Error bar represents standard error of the mean (SEM). Statistically significant differences between groups are reported in Table S7 using Mann-Whitney U test. Gating strategy is shown in Supplementary Fig. S1A–C. Source data are provided as a Source Data file. **C** Heatmap of normalised values demonstrate increased numbers of cells in splenic tissue at earlier timepoints (4 – 6 weeks after infection) followed by increased numbers of cells in draining lymph node (DLN) at later timepoints (10 and > 10 weeks) after infection (C57BL/6 mice controls: week (W) 4 – 6 n=5 per group, W10 n=3, W>10 n=12; ∼10 CFU dose: W4 n=4, W6 – 10 n=5 per group, W>10 n=9; ∼100 CFU dose: W4 – 10 n=5 per group, W>10 n=10).

There were also significant differences in innate immune cell numbers in C57BL/6 mice. In splenic tissue 4 – 6 weeks after low dose challenge, compared to control mice, there were increased levels of γδ T cells, neutrophils, macrophages (all p=0.016) and dendritic cells (p=0.032) (Fig 3B; Table S7). Following high dose challenge, differences in innate splenic immune responses were not as apparent and, in some cases, were again significantly below values observed in the low dose group, including γδ T cells, macrophages (both p=0.016) and dendritic cells (p=0.032) at 4 weeks (Table S7).

In the interval 6 – 10 weeks after challenge, there were no significant differences between the two C57BL/6 dosing groups. In mice that had reached the humane end point (*i.e*. those culled > 10 weeks after challenge), there were significantly more macrophages (p=<0.001) and dendritic cells (p=0.012) in DLNs of mice challenged with a low dose compared to the high dose, and numerous adaptive and innate immune features were significantly greater in the low-dose model compared to controls (Table S7).

With multiple immune features compared over various timepoints, adjustment for repeated measures was performed to understand which comparisons were most unlikely to be detected by chance alone, including markers of activation. This false discovery adjustment highlighted the significance of adaptive cells, particularly CD8^+^ cells, in C57BL/6 responses to *M. ulcerans* low-dose infection. In C57BL/6 mice with ulcerative disease, activated CD38^+^CD8^+^ cells were significantly more numerous in DLNs of low dose mice compared to both controls and high dose mice (Table S8).

When immune cellular values were normalised and compared over time, an early splenic increase in cell numbers, followed by later increases in DLNs, were most apparent in the C57BL/6 low dose model (Fig 3C). The blunted cellular immune responses observed in the high dose infected mice might be explained by presumed higher mycolactone concentrations associated with the increased bacterial burden.

### BALB/c cellular responses to infection are not dose-dependent within the dose range tested

For BALB/c mice, there were no significant differences between low dose and high dose mice in either adaptive (Fig 4A) or innate (Fig 4B) immune profiles at any timepoint. Unlike C57BL/6 mice, splenic cellular values in BALB/c mice were generally similar to controls, or lower than observed in control mice, particularly 10 weeks following challenge in both high and low dose groups (Table S7). When values were normalised and analysed over time; again, there was no clear pattern to emerge (Fig 4C). These results suggest that a robust cellular response, as observed in C57BL/6 mice, may be an important factor in increased disease pathology, and poorer clinical outcomes.

**Figure 4.**
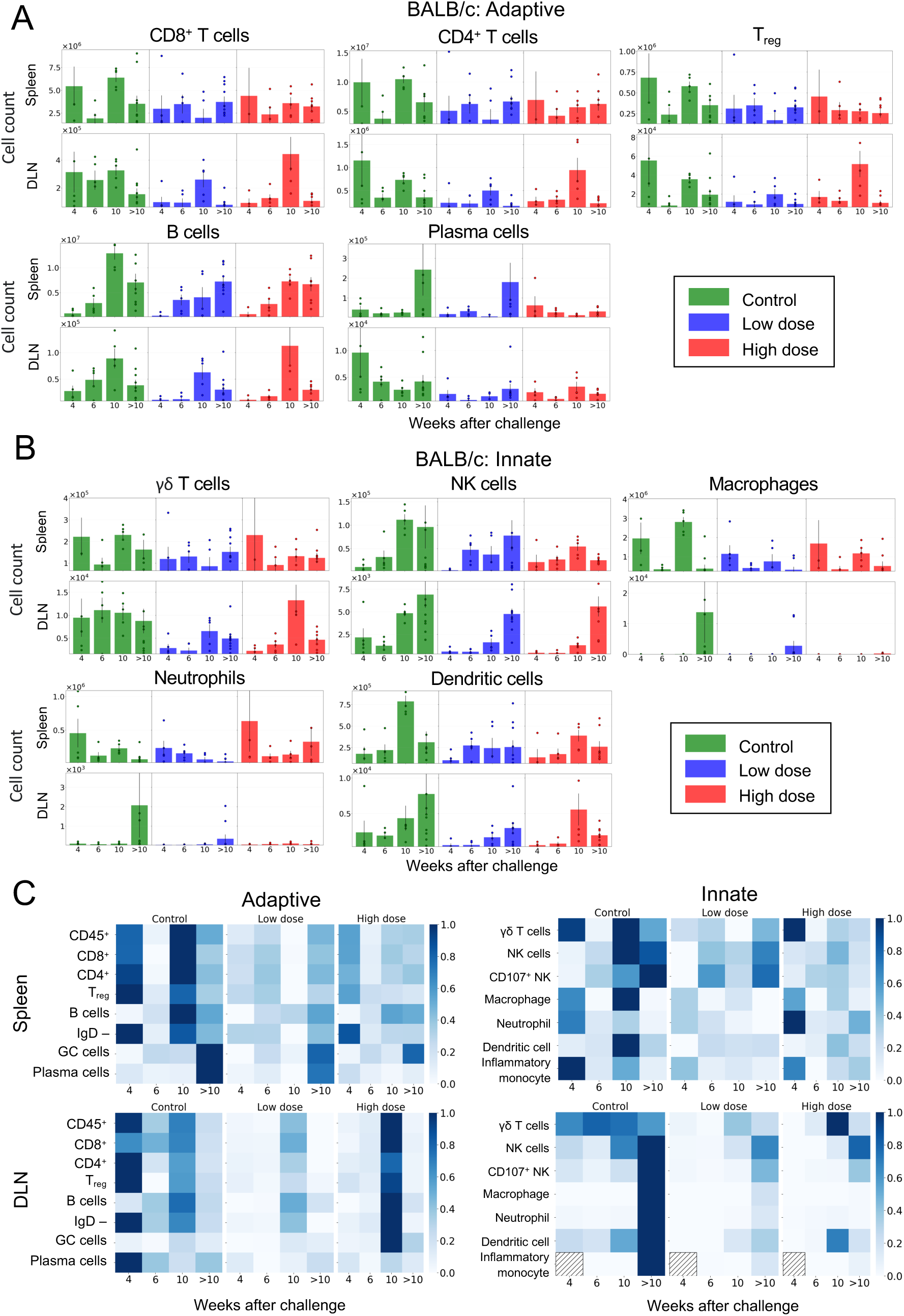
Total cell counts of adaptive and innate immune features over time in BALB/c mice following low-dose *M. ulcerans* infection. **A** Adaptive and **B** innate immune features over a maximum of 19 weeks using absolute numbers of effector CD8^+^ T cells (CD8^+^ CD62L^lo^ CD44^hi^), effector CD4^+^ T cells (CD4^+^ CD62L^lo^ CD44^hi^), T_reg_ cells (CD4^+^ CD25^+^ CD62L^+^), B cells (CD19^+^ B220+), plasma cells [antibody secreting cells] (IgD-B220^lo^ CD138+), γδ T cells (TCRγδ CD3+), NK cells (CD49b+ NK1.1+), macrophages (CD64^+^ F4/80^+^), neutrophils (CD11b^+^ Ly6G^+^) and dendritic cells (CD11c^+^MHCII^+^). Error bar represents SEM. Statistically significant differences between groups are reported in Table S7 using Mann-Whitney U test. Gating strategy is shown in Supplementary Fig. S1A–C. Source data are provided as a Source Data file. These are visualised as a heatmap **C** of normalised values, which do not demonstrate any clear pattern over time (BALB/c mice controls: week (W) 4 – 6 n=5 per group, W10 n=3, W>10 n=10; ∼10 CFU dose: W4 – 10 n=5 per group, W>10 n=10; ∼100 CFU dose: W4 n=4, W6 – 10 n=5 per group, W>10 n=8).

### High dose challenge is associated with detection of serum anti-M. ulcerans antibodies

To investigate humoral immune responses induced by infection, serum from infected C57BL/6 and BALB/c mice were analysed for antibody responses against *M. ulcerans* whole cell lysate by ELISA. In both C57BL/6 and BALB/c mice infected with the low dose, there were no significant differences in antibody titres compared to control mice at each time point. In comparison, high dose C57BL/6 mice that progressed to the endpoint of ulceration (>10 weeks) had significantly higher anti-*M. ulcerans* antibody titres than low dose mice (Fig 5A). High dose BALB/c mice also had higher antibody titres at the endpoint of ulceration compared to low dose and control mice (Fig 5B). These data suggest that in mice antibody responses to *M. ulcerans* are observed late in the course of infection, perhaps explained by increased extra-cellular bacteria at the later infection timepoints.

**Figure 5.**
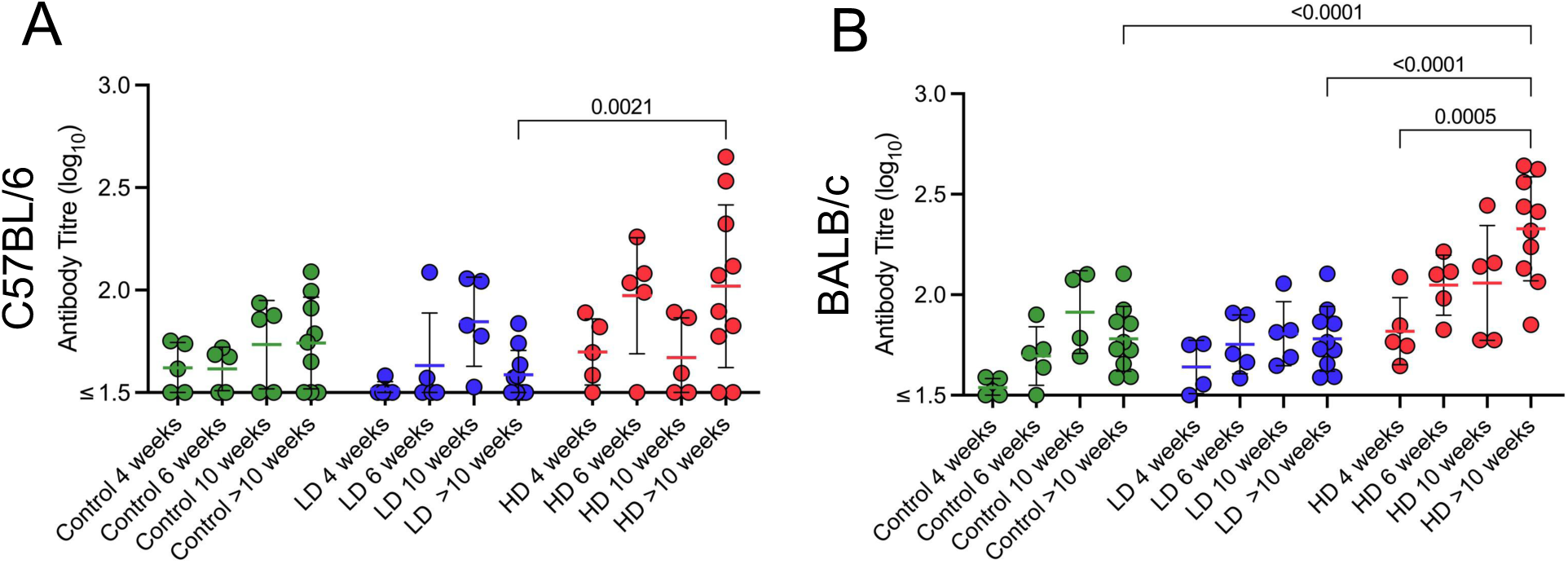
Antibody responses to low dose (‘LD’) and high dose (‘HD’) infection over time. **A** Antibody titres in C57BL/6 and **B** BALB/c serum over maximum 19 weeks (BALB/c) or maximum 15 weeks (C57BL/6) were measured by ELISA against bacterial whole cell lysate. (C57BL/6 controls: Week (W) 4 – 10 n=5 per group, W>10 n=9; C57BL/6 both dosing groups: W4 – 10 n=5 per group, W>10 n=10; BALB/c controls: W4 – 6 n=5 per group, W10 n=4, W>10 n=10; BALB/c ∼10 CFU: W4 n=4, W6 – 10 n=5 per group, W>10 n=10; BALB/c ∼100 CFU: W4 – 10 n=5 per group, W>10 n=10). Statistical differences were determined using two-way ANOVA. Error bars represent the SEM.

### Inflammatory responses to low dose M. ulcerans in C57BL/6 mice are dominated by activated adaptive immune responses, which are associated with delayed disease progression

To define immune features associated with each respective dosing group, Random Forest classifiers were first employed to evaluate the combined ability of immune parameters and their activation markers (features) to discriminate between low and high dose challenge groups. This analysis identified the top five immune parameters contributing to group separation for each mouse strain, highlighting key features driving the classification. In C57BL/6 mice (Fig 6A), this analysis highlighted the features that might be driving the inflammatory responses observed in the low dose C57BL/6 model; most notably, DLN dendritic cells and CD107^+^ NK cells, and activated KLRG1^+^ and CD25^+^ splenic CD8^+^ cells. In BALB/c mice (Fig 6B), activated (germinal centre) DLN B cells emerged as the top predictive feature of inoculation dose. Classifier performance was evaluated by comparing the area under the curve (AUC) from the actual data to that from 100 repetitions with randomised class labels (Fig 6C). The difference in AUC between the actual data and the randomised runs was minimal for the BALB/c group, while a significant difference, exceeding two standard deviations from the randomised mean, was observed for the C57BL/6 mice. These data confirm that robust cellular immune responses occur in C57BL/6 mice during *M. ulcerans* infection, but not in BALB/c mice. Correlation analyses of clinical and immune features confirmed these patterns (Fig S4C).

**Figure 6.**
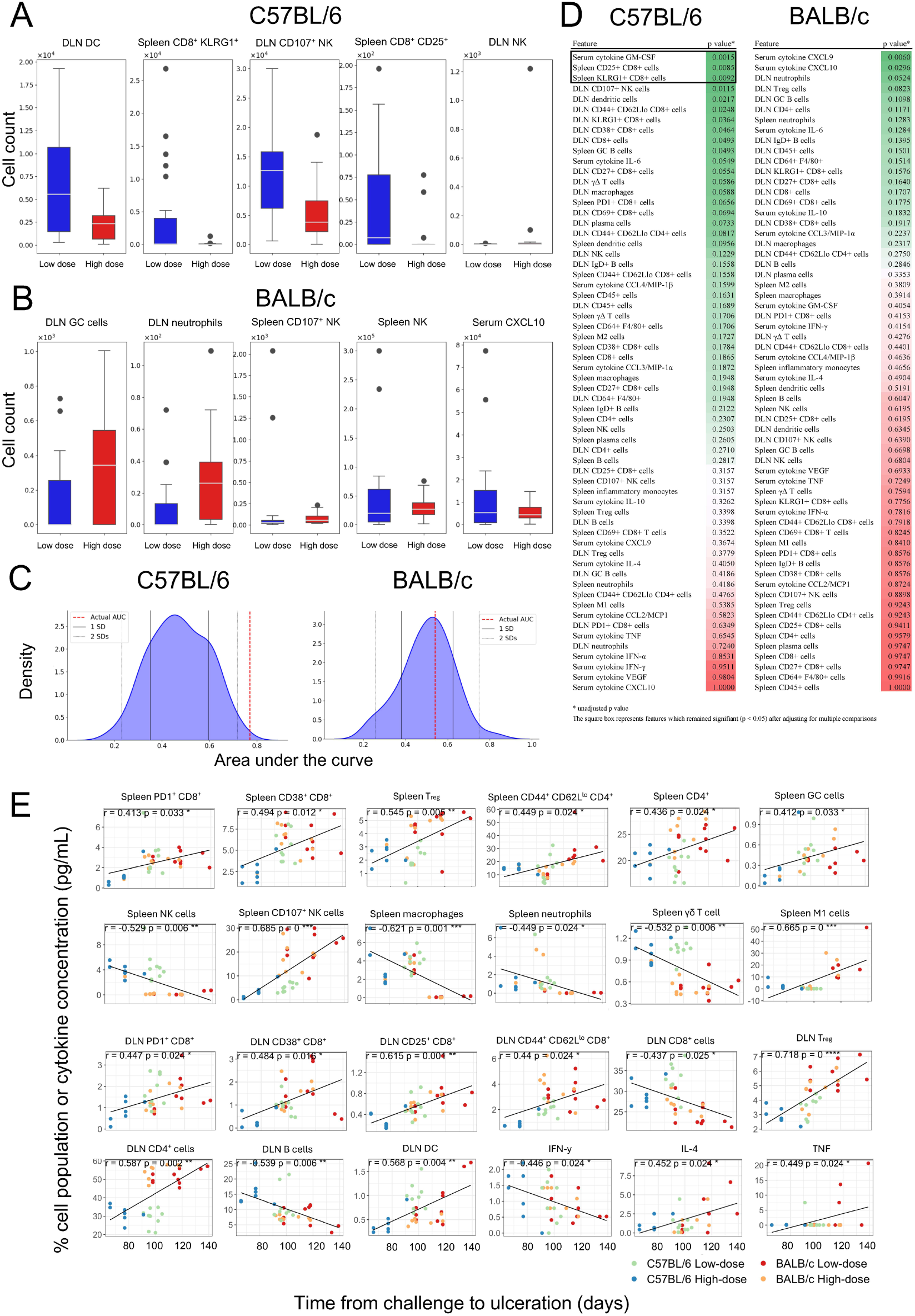
Predictive features associated with challenge ‘dose’. Figures **A** and **B** demonstrate the five features ranked by importance using random forest classifiers; error bars represent SEM. **C** shows classifier performance demonstrated by the difference in AUC between models trained on actual data and those using randomised labels. **D** demonstrates the significance between high dose and low dose groups using univariable Mann-Whitney U tests, with p-value reported alongside each immunological parameter; the square box represents features which remained significant after adjusting for multiple comparisons, To adjust for false discovery due to multiple comparisons, from each group of highly correlated variables (correlation coefficient > 0.8), only one representative variable was retained, and the Benjamini–Hochberg procedure was then applied; these values are also displayed in Table S9. **E** shows the correlation between various immune features and the time to ulceration, with statistical significance of the Pearson correlation coefficient (r) determined using Student’s t-tests, adjusting p values for multiple comparisons using the Benjamini-Hochberg (false discovery rate) method. (C57BL/6: ∼100 CFU n=7, ∼10 CFU n=9, BALB/c: ∼100 CFU n=8, ∼10 CFU n=10).

Univariate tests were also performed to evaluate differences in individual immune features between the dosing groups over the entire experimental time course (Fig 6D). Several significant differences were observed in C57BL/6 mice. These differences were predominantly among CD8^+^ T cells and associated responses and included splenic CD25^+^ and KLRG1^+^ CD8^+^ T cells (p = 0.036) and serum cytokine GM-CSF (p = 0.020) (Table S9). No differences in immune features were observed for BALB/c mice between dose groups at any time point.

Finally, to assess whether any immunological features were positively or negatively correlated with disease progression, immune features were analysed for each mouse strain across both doses over time (Fig 6E). In splenic tissue, increased frequencies of immune cells correlating with delayed disease progression included activated (CD107^+^) NK cells, CD4^+^ cells, CD4^+^ effector (CD44^+^CD62L^lo^) cells, regulatory T cells, as well as activated (germinal centre) B cells and CD38^+^ and PD1^+^ CD8^+^ cells. The increased splenic immune cells that correlated with more rapid disease progression included macrophages, neutrophils and γδ T cells. In DLNs, a correlation with delayed disease progression was observed with increased presence of dendritic cells, CD4^+^ cells, regulatory T cells as well as PD1^+^, CD38^+^ and CD25^+^ CD8^+^ cells, and effector (CD44^+^ CD62L^lo^) CD8^+^ cells. Of note, increased total DLN CD8^+^ cell frequencies correlated with more rapid disease progression, underscoring the importance of CD8^+^ activation in delaying disease progression. Notably, although splenic macrophages were associated with disease progression, M1-biased splenic macrophages correlated with delayed disease progression. In this study, the cytokine IFN-γ was correlated with more rapid disease progression, likely influenced by the greater IFN-γ responses in mice that received the higher challenge dose, suggesting dysregulated immune responses. These analyses demonstrate the important role of adaptive immunity, and particularly activated T cells, following low dose *M. ulcerans* infection.

## Discussion

The principal goal of this study was to demonstrate the importance of accurately dosing *M. ulcerans* with a realistic bacterial inoculum and to establish a relevant animal model of BU to characterise host-pathogen responses. In Australia, *Aedes notoscriptus* mosquitos are a vector of *M. ulcerans*, harbouring a mean of ∼300 total *M. ulcerans* bacteria per mosquito (range, 11 – 4,200 bacteria), as established by qPCR [21]. The actual number of viable bacteria transmitted during mosquito blood-feeding is likely to be even lower than total bacterial estimate. Here, a clear dose-response relationship in both BALB/c and C57BL/6 mice was demonstrated, with low doses extending the time to lesion onset, replicating the long incubation periods seen in humans [5,6]. Mice challenged with a lower dose also demonstrated slower progression of clinical disease, which is an important safety consideration if such dose ranges are to be used for a human challenge study, as more rapidly progressive disease is an unfavourable outcome [22,23]. Ideally, in a controlled human infection, lesions would progress slowly as is typical in naturally occurring BU, and slower disease progression will also enable timely therapeutic intervention.

With a 100% attack rate in both groups of mice at both low and high doses, this study confirms the highly infectious nature of *M. ulcerans* in susceptible hosts. This study should encourage researchers to reframe the concept of a ‘high dose’ *M. ulcerans* challenge and highlights the importance of host factors on BU clinical outcomes. Compared to BALB/c mice, C57BL/6 mice demonstrated a more oedematous/inflammatory phenotype, as evidenced clinically and upon detailed immunological analysis. Injecting additional *M. ulcerans* extracellular matrix alongside the bacterial inoculum may in itself trigger inflammatory response [24]; the present study avoided that possibility, by using a carefully prepared suspension of individual bacilli, with clumps filtered prior to cell banking [13].

The natural history of *M. ulcerans* infection involves a cycle of infection, intracellular bacillary replication, leukocyte destruction and extracellular propagation [25]. At least in C57BL/6 mice, we found that the systemic activation of cellular responses is only apparent in mice challenged with low *M. ulcerans* doses. Although the implications of this require further research, it supports the hypothesis that immune responses to vaccination may be ‘overwhelmed’ by unrealistically high challenge doses. This will require further interrogation with promising candidate vaccinations [9] prior to low dose challenge, particularly in C57BL/6 mice.

Reinforcing the importance of host-specific immunity, previous research demonstrated the markedly pronounced cellular response to *M. ulcerans* infection in the DLNs of C57BL/6 mice compared to BALB/c mice [24]. Our study also suggests that a systemic cellular response occurs soon (∼1 month) after low dose infection, presumably promoting the recruitment of cells from the systemic circulation to the site of infection. Considering the early intracellular phase of *M. ulcerans* infection [17], it is likely that the Th1-bias of this mouse strain enables the swift recruitment of macrophages, thereby enabling a more rapid infection and inflammatory clinical phenotype. It has been assumed that phagocytes play a key role in early host defense against *M. ulcerans* [26]. Counterintuitively, a strong phagocytic response may act to expose mycobacteria to susceptible cells [17], potentially enabling rapid uptake, replication, and propagation of infection. Indeed, the present study does suggest a correlation between increased systemic (*i.e.* early) phagocytic responses and more rapid progression of clinical disease. Co-opting the immune system is also known to occur in *M. marinum* infection, a related mycobacterium with close genetic homology [27]. Intra-phagocyte trafficking of *M. ulcerans* to DLNs is observed in C57BL/6 [28] and BALB/c [29] mice and may be more likely to occur early in the course of disease, when mycolactone concentration is low; nevertheless, due to the organism’s restricted temperature range, disease in organs might be contained by the host’s immunological response to non-viable bacilli. In the natural animal host, the native Australian possum, evidence of dissemination and cell-mediated immune control has been reported using experimental infection with *M. ulcerans* JKD8049 [30]. A limitation of the present study is that we did not collect microbiological data to demonstrate systemic dissemination of infection. Previous reports have been unable to demonstrate the presence of *M. ulcerans* in the spleens of C57BL/6 mice [28], thus the mechanism of systemic immune stimulation in these mice remains unclear. Nevertheless, here we have shown that a robust, early, systemic, Th1-biased cellular immune response is unable to protect C57BL/6 mice from challenge with a low number of *M. ulcerans* bacilli.

This study demonstrated the important role of specific cellular responses to low dose *M. ulcerans* infection. In the low dose C57BL/6 model, it appears that early splenic cellular responses are broad, including both innate and adaptive immune cells. Univariate and machine learning analyses showed that immune responses to low dose infection are dominated by activated CD8^+^ T cell responses and correlation models show that activated CD8^+^ T cells in spleen and DLN are associated with a delayed disease course (Fig 6). However, the specific mechanisms by which CD8^+^ T cells fail to protect from ulceration remain to be understood. Investigating the mechanism(s) of this failure has potentially important implications for creating effective vaccination strategies, such as building vaccines that improve antigenic presentation and stimulate *M. ulcerans-*specific CD8^+^ T cell responses. For instance, saponin-based adjuvants strongly induce cross-presentation by dendritic cells, leading to subsequent CD8^+^ T cell activation [31,32]. Using such adjuvants with promising subunit vaccine candidates may offer additional protection from infection via antigen-specific CD8^+^ T cell responses and warrants consideration in future research.

Dendritic cells also have a prominent role in low dose *M. ulcerans* infection, with more dendritic cells observed in the DLNs of C57BL/6 mice later in the course of disease. DLN dendritic cells appeared significant in univariate analysis, although this difference just exceeded the statistical significance threshold when correcting for multiple comparisons (p=0.050). Dendritic cells were the top classifier feature explaining the difference between the high and low dose models. Finally, DLN dendritic cells were also associated with delayed disease progression, highlighting the important role of these key antigen presenting cells in low dose *M. ulcerans* infection.

Our data suggest that a systemic reduction in immune cellularity occurs in C57BL/6 mice as bacterial burden increases over time (Fig 2A). This phenomenon was not observed in our study in BALB/c mice, nor in previous research [29]. It remains unclear whether this reduction reflects true systemic immune suppression in C57BL/6 mice, or regional sequestration of cells at the site of infection. Considering the remarkably oedematous/inflammatory nature of the infection locoregionally, the latter is favoured to be the likely mechanism. As reported by Converse and colleagues [33], we found that *M. ulcerans* growth kinetics were similar in both BALB/c and C57BL/6 mice across all timepoints, so any differences observed between each mouse strain are unlikely to be influenced by host control of bacterial growth or differential mycolactone concentration. Nevertheless, a limitation of this study is that mycolactone concentrations were not measured, therefore we cannot exclude the influence of potential variable mycolactone expression over time or between each mouse strain.

Another limitation of this research is that very early immune responses (< 4 weeks) were not captured. Oliveira *et al*. previously demonstrated acute neutrophilic infiltration at the site of inoculation in BALB/c mice, although their study used a very large inocula (1 x 10^5^ AFB) of a virulent African (mycolactone A/B-producing) isolate which resulted in leukocyte destruction as early as 24 hours after inoculation [25]. The authors also showed that inoculation with ∼ 316 AFB had a similar histopathological outcome with delayed kinetics, which was reinforced by the present study. Nevertheless, neutrophils were not observed in tail histology at any of our earliest timepoints. A previous BALB/c mouse model of infection by Fraga *et al.* demonstrated early accumulation of CD4^+^ lymphocytes in DLN tissue (around 14 days post challenge). Their study is difficult to compare and may be less generalisable, as ulceration occurred within 24 days of challenge, compared to an average of 116 days in BALB/c mice in our low dose challenge model. They also used a very high inoculation dose of a virulent African isolate (3 x 10^5^ AFB) using a footpad site of inoculation [29].

Although the murine footpad model of BU has been utilised extensively in previous vaccine research [9], the footpad model has numerous limitations. Footpad infection more readily exposes lesions to the environment, which may increase the risk of irritation and superinfection, confounding markers of inflammation. Similar to the footpad site, the tail model shares the convenience of little hair coverage. The tail also appears a favoured site for the development of BU lesions, even in mice inoculated intracerebrally [34]. As the foot is a weightbearing structure, mice may develop gait abnormalities, and although pain is uncharacteristic for this disease, the confined anatomical space in the footpad may predispose mice to discomfort associated with tissue swelling. Previous studies have identified the hock as an ethically acceptable alternative to the footpad model [24], although the tail was not specifically characterised, and the high *M. ulcerans* dose (∼10^7^ CFU) was based on the wet-weight of injected material, which is inherently less accurate than the enumeration method reported in the present study. An additional practical benefit of the tail model is that it is easier to inspect during routine observation, with less handling required when compared to sites located over the limb. The tail is also a reproducible site of infection, as the incubation period in naïve BALB/c mice receiving a low dose challenge in the present study was identical to our initial pilot model using *M. ulcerans* JKD8049 from the same cell bank used in the present study [15]. The refined tail model presented here today therefore represents an ethically acceptable, reproducible and convenient site for future murine BU research.

Although we used BALB/c and C57BL/6 mice to understand the infection rate in mammalian hosts with broad immune biases, we are unable to predict whether this will similarly translate to a high infection rate in a human model of infection. Similar to our study, Coutanceau *et al*. did not observe granuloma formation in either BALB/c or C57BL/6 mice [28], which is unlike human cases, where granulomas are associated with healing [14]. Genetic differences between in-bred laboratory mice and humans are likely to confer varying degrees of susceptibility to infection. For example, both BALB/c and C57BL/6 mice are known to contain polymorphisms in the natural resistance-associated macrophage protein gene (NRAMP1) [35]; in humans, polymorphisms in this gene are also associated with increased susceptibility to *M. ulcerans* infection, although the majority of individuals selected for challenge are unlikely to carry these polymorphisms [36].

## Conclusion

By analysing an infection time-course of over 70 clinical, immunological and microbiological parameters, this study identified dose-related differences in outcomes following *M. ulcerans* infection in mice. We observed that strong cell-mediated responses were associated with more rapid Buruli ulcer onset. In C57BL/6 mice, low bacterial challenge doses trigger an early systemic immune response, well before the onset of clinical disease, enabling the trafficking of immune cells to the site of infection while the infection develops. In the higher dose challenge model, these immunological features were likely masked by higher bacterial concentrations and (presumably) more rapid mycolactone accumulation. Antibody responses were not evident following low dose infection, and antibody presence after high dose infection did not predict disease protection. Given these findings and considering recent evidence suggesting that low inocula are likely acquired during natural infection, we suggest that future pre-clinical research considers using low challenge doses. In the context of a planned controlled human infection of *M. ulcerans*, this study supports lower challenge doses that will slow the course of clinical disease. A reliable, slowly progressing infection is preferred to a rapid-onset disease phenotype. Vaccination approaches that facilitate CD8^+^ T cell activation in response to *M. ulcerans* infection may facilitate protection from disease.

## Acknowledgements

We thank Melbourne Histology Platform for preparation of histology and Phenomics Australia for histology reporting. We acknowledge the Melbourne Cytometry Platform (The Doherty Institute) for provision of flow cytometry services. We also thank Melbourne Bioresources Platform staff at the Doherty Institute.

## Author contributions

TPS, KK, AHB and BYC supervised the study. SM, IJHF, LK, BYC designed the experiments. SM, IJHF, AHB, JLP, BH, LK, BYC performed and analysed experiments. SM, IJHF, LK, AHB, HAM and BYC analysed data. SM, IJHF, LK, KK, AHB, BYC and TPS wrote the manuscript. All authors reviewed and approved the manuscript.

## Data availability

All primary data not included in the main study or supplementary files are available here: https://figshare.com/s/2c9befed57e953f61a63.

## Supplementary material

**Figure S1.**
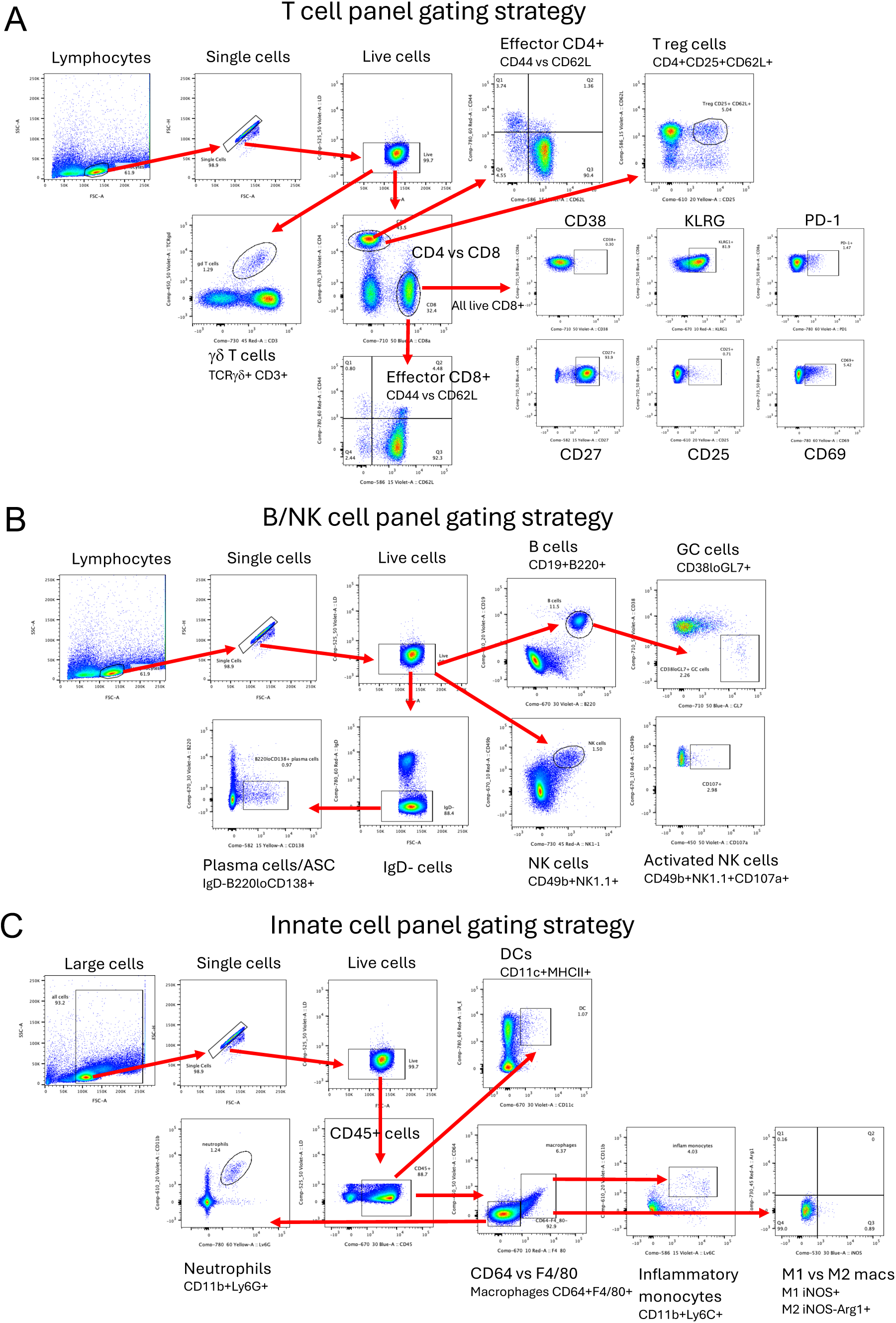
Gating strategy for flow cytometry. **A** T cells, **B** B/NK cells and **C** innate immune cells.

**Figure S2.**
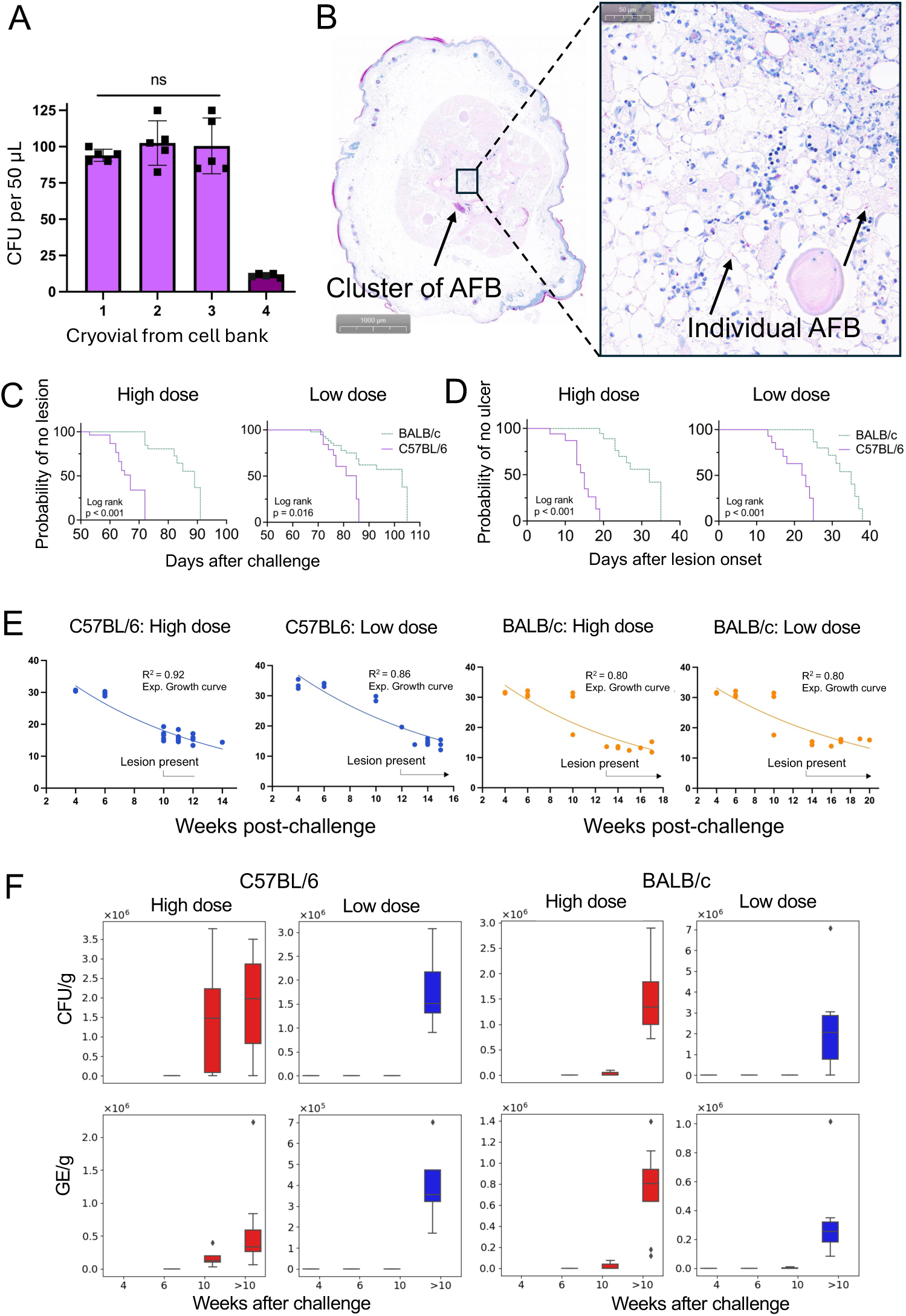
Microbiological and clinical features over time. **A** CFU challenge dose enumeration as calculated from each vial prior to inoculation; the three columns on the left show the CFU count in the vials used for high dose challenge. The column on the far right illustrates the mean CFU value injected after dilution to a low dose. Errors bars represent 95% confidence interval. **B** Cross section through the ulcerated tail of a C57BL/6 mouse (#HB8.5) challenged with ∼ 100 CFU of *M. ulcerans* JKD8049 and stained with Ziehl–Neelsen stain; there is a large and dense clump of *M. ulcerans* acid fast bacilli (AFB) immediately adjacent to the vertebral bone; inset demonstrates individual AFB visible within the bone marrow tissue (inset bar represents 50 μm length at magnification of 20x). **C** survival curves demonstrating the time to lesion onset in C57BL/6 mice compared to BALB/c mice (∼10 CFU challenge dose: C57BL6 n=10, BALB/c n=10; ∼100 CFU challenge dose: C57BL6 n=15, BALB/c n=10). **D** survival curves demonstrating the time from lesion onset to ulceration in C57BL/6 mice compared to BALB/c mice (∼10 CFU challenge dose: C57BL6 n=10, BALB/c n=10; ∼100 CFU challenge dose: C57BL6 n=12, BALB/c n=10). **E** visualises *M. ulcerans* IS*2404* PCR cycle threshold over time according to dose and mouse type (∼10 CFU dose C57BL/6: week (W) 4 n=3, W6 n=3, W10 n=3, W>10 n=10; ∼100 CFU dose, C57BL/6: W4 n=3, W6 n=3, W10 n=5, W>10 n=10; ∼10 CFU dose, BALB/c: W4 n=3, W6 n=3, W10 n=3, W>10 n=10; ∼100 CFU BALB/c: W4 n=3, W6 n=3, W10 n=3, W>10 n=10). The line represents the exponential growth curve. R^2^ indicates the goodness-of-fit. **F** microbiological enumeration (in CFU/g and GE/g) of bacilli over time in both low and high dose cohorts, error bars represents SEM.

**Figure S3.**
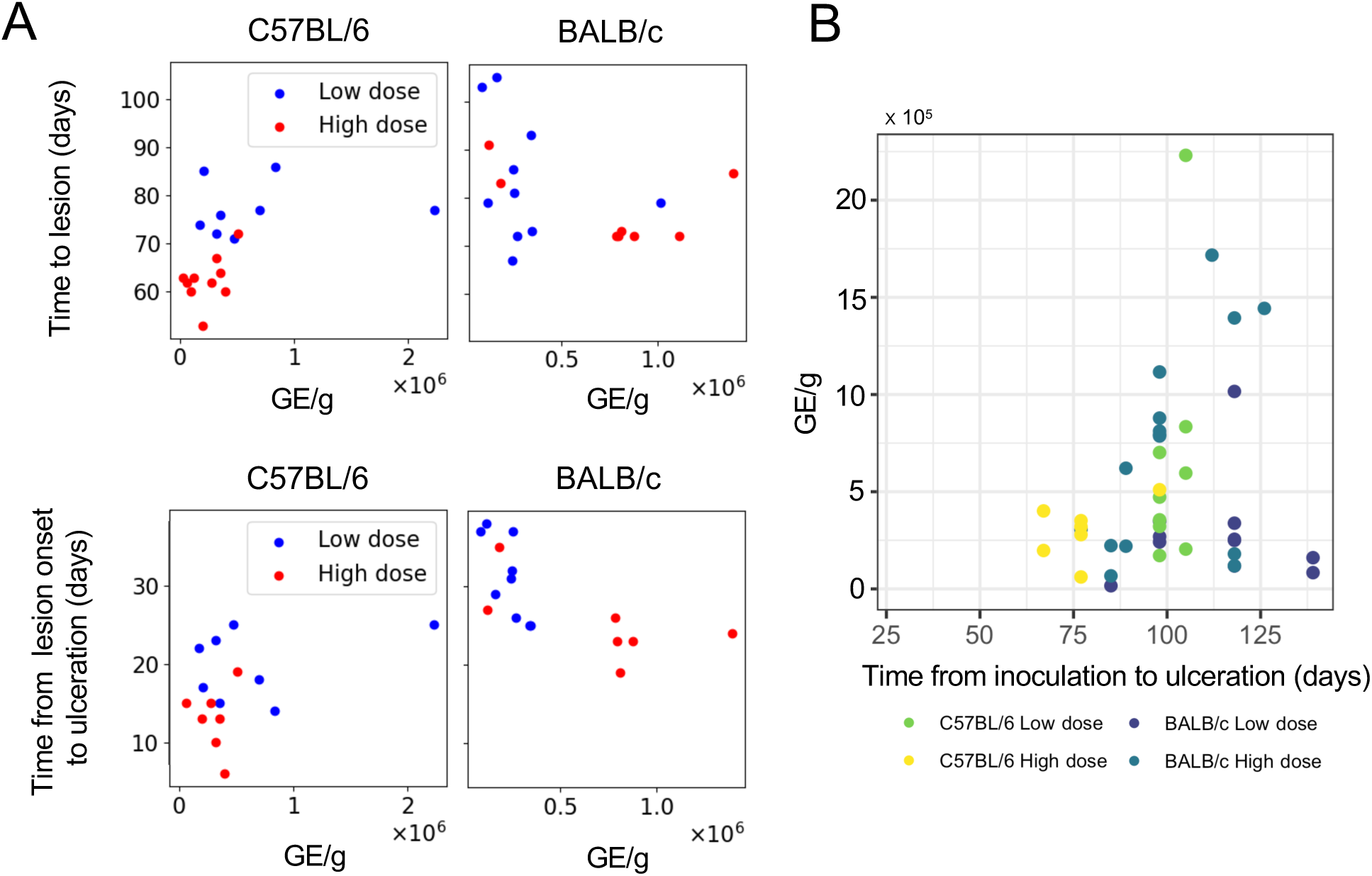
Microbiological correlates of clinical disease over time. **A** the relationship between clinical features of infection (time to lesion onset and time from lesion onset to ulceration) and correlation with the burden of bacilli in GE/g. **B** correlation plot between GE/g and time from inoculation to ulceration across both mouse lines and dosing groups.

**Figure S4.**
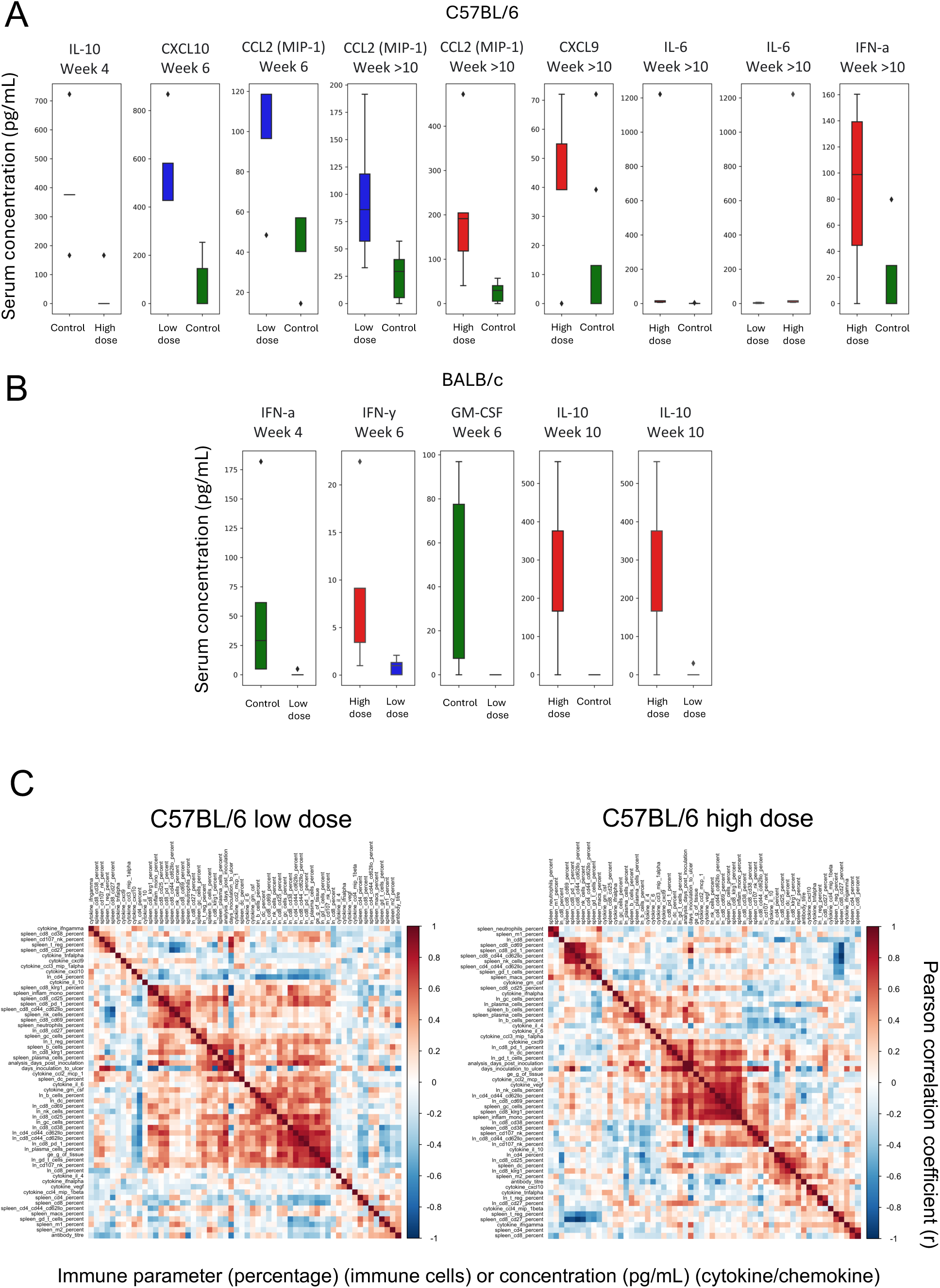
Statistically significant differences in cytokines and chemokines at various experimental timepoints in **A** C57BL/6 mice and **B** BALB/c mice; error bar represents the SEM. **C** correlation matrix of clinical and immune features over time; dark red (Pearson correlation coefficient = 1) suggests a strong correlation, dark blue (Pearson correlation coefficient = -1) suggests a strong negative correlation.

**Table S1.** Clinical, microbiological and histopathological features: C57BL6 mice, high dose.

| Time after challenge (weeks) | Mean IS2404 PCR Ct | GE /g of tissue | Lesion | Lesion type | CFU/g tissue | Histology report |
| --- | --- | --- | --- | --- | --- | --- |
| 4 | N/A | N/A | No | N/A | N/A | No lesions seen. |
| 4 | N/A | N/A | No | N/A | N/A | No lesions seen. |
| 4 | 30.73 | 84.53 | No | N/A | No growth | N/A |
| 4 | 30.73 | 65.75 | No | N/A | No growth | N/A |
| 4 | 30.37 | 98.43 | No | N/A | No growth | N/A |
| 6 | N/A | N/A | No | N/A | N/A | No lesions seen. |
| 6 | N/A | N/A | No | N/A | N/A | No lesions seen. |
| 6 | 30.28 | 77.66 | No | N/A | No growth | N/A |
| 6 | 29.63 | 100.11 | No | N/A | 152.0 | N/A |
| 6 | 28.85 | 206.15 | No | N/A | 119.00 | N/A |
| 10 | 17.13 | 1.01E+05 | Yes | Pre-ulcerative | 1.47E+06 | Multifocal dermal and subcutaneous oedema, necrosis, moderate macrophage and lymphocyte infiltrations. Necrosis associated with AFB. |
| 10 | 14.81 | 4.01E+05 | Yes | Ulcer | 2.23E+06 | Mild dermal and moderate subcutaneous oedema, mild necrosis, infiltrations of macrophages and lymphocytes. AFB present in one area. |
| 10 | 19.28 | 2.68E+04 | Yes | Pre-ulcerative | No growth | Moderate to severe multifocal to diffuse dermal and subcutaneous oedema with scanty macrophages (occasionally with scanty AFB in the cytoplasm) and neutrophil infiltration. Destruction of blood vessel walls and skeletal muscle and tendon necrosis, areas associated with multifocal abundant acid-fast bacilli. No epidermal necrosis. |
| 10 | 16.63 | 1.25E+05 | Yes | Pre-ulcerative | 7.88E+04 | Moderate multifocal to diffuse dermal and subcutaneous oedema, vasculopathy and infiltration of macrophages, lymphocytes, neutrophils. Multifocal skeletal myofiber necrosis with multifocal AFB. |
| 10 | 15.55 | 1.97E+05 | Yes | Ulcer | 3.76E+06 | Severe multifocal full-thickness epidermal necrosis, multifocal epidermal pustule formation and diffuse subcutaneous oedema, with infiltrations of macrophages, lymphocytes, neutrophils. There is vasculitis, and tissue necrosis including skeletal myofiber necrosis with multifocal AFB. |
| 11 | 16.11 | 3.06E+05 | Yes | Ulcer | 5.67E+05 | Severe locally extensive full-thickness epidermal necrosis, multifocal epidermal vesicles, diffuse subcutaneous oedema, multifocal vasculopathy, infiltrations of macrophages, lymphocytes, neutrophils. Multifocal necrosis of connective tissue and skeletal muscle with multifocal AFB. |
| 11 | 18.37 | 6.14E+04 | Yes | Ulcer | 2.17E+06 | Severe locally extensive full-thickness epidermal necrosis, diffuse dermal and subcutaneous oedema, multifocal vasculopathy, infiltrations of macrophages, lymphocytes, neutrophils. Multifocal necrosis of connective tissue, skeletal muscle and bone marrow with multifocal AFB. |
| 11 | 15.50 | 3.51E+05 | Yes | Ulcer | 1.10E+06 | Severe locally extensive full-thickness necrosis of the epidermis. The intact epidermis shows multifocal hyperplasia and overlies markedly oedematous dermis and subcutaneous with infiltrations of inflammatory cells, macrophages, lymphocytes, neutrophils; tissue necrosis, vasculopathy and necrosis of skeletal muscles. AFB are scattered throughout the oedematous and necrotic tissue in low numbers. |
| 11 | 15.08 | 2.79E+05 | Yes | Ulcer | 1.77E+06 | Severe locally extensive full-thickness necrosis of the epidermis. The intact epidermis shows multifocal hyperplasia. Severe multifocal to coalescing dermal and subcutaneous oedema with infiltrations of inflammatory cells, macrophages, lymphocytes, neutrophils; tissue necrosis, vasculopathy and necrosis of skeletal muscles. AFB are present scattered and within necrotic tissue. |
| 11 | 14.54 | 3.20E+05 | Yes | Ulcer | 3.50E+06 | Focal epidermal erosion overlying an area of mild subcutaneous oedema, inflammation with scanty acid-fast bacilli. Moderate to severe extensive dermal and subcutaneous oedema, multifocal vasculopathy, infiltrations of macrophages, lymphocytes, neutrophils. Multifocal necrosis of connective tissue and skeletal muscle with multifocal AFB. |
| 12 | 15.27 | 2.23E+05 | Yes | Ulcer | 4.45E+05 | Focal epidermal erosion, multifocal hyperplasia and diffuse hyperkeratosis, severe segmental subcutaneous oedema, mixed inflammatory cell infiltrates multifocal necrosis of skeletal muscle and bone marrow, and focal vasculitis. AFB associated with necrotic tissue and within the wall of the inflamed blood vessels. |
| 12 | 15.83 | 2.20E+05 | Yes | Ulcer | 1.35E+06 | Multifocal full thickness epidermal necrosis. Moderate diffuse subcutaneous oedema with macrophage, neutrophils, lymphocytes, multifocal tissue necrosis, skeletal muscle and into the bone marrow. Multifocal AFB associated with necrosis and in the bone marrow. |
| 12 | 13.39 | 6.21E+05 | Yes | Ulcer | 2.66E+06 | Severe diffuse subcutaneous oedema, inflammatory infiltrates (macrophages, lymphocytes, neutrophils), vasculopathy, tissue necrosis with AFB. Multifocal epidermal hyperplasia, hyperkeratosis and focal pustule formation. |
| 12 | 17.09 | 6.71E+04 | Yes | Ulcer | 1.54E+04 | Moderate to severe multifocal to diffuse dermal and subcutaneous oedema with scattered macrophages, lymphocytes and neutrophils associated with multifocal AFB. Focal vasculopathy. |
| 14 | 14.38 | 5.10E+05 | Yes | Ulcer | 2.83E+06 | Severe full thickness skin and subcutaneous necrosis of >60% of the section with deep ulceration and loss of adnexal structures. Intense inflammatory infiltrates of neutrophils, macrophages, lymphocytes are present. Severe vasculopathy associated with connective tissue and skeletal muscle necrosis. Severe subcutaneous oedema of the remaining section with multifocal necrosis and skeletal muscle necrosis and moderate mixed inflammatory infiltrates. Multiple clusters of AFB in affected areas. |

**Table S2.** Clinical, microbiological and histopathological features: C57BL6 mice, low dose.

| Time after challenge (weeks) | Mean IS2404 PCR Ct | GE /g of tissue | Lesion | Lesion type | CFU/g tissue | Histology report |
| --- | --- | --- | --- | --- | --- | --- |
| 4 | N/A | N/A | N/A | N/A | N/A | N/A |
| 4 | N/A | N/A | No | N/A | N/A | No lesions seen. |
| 4 | 33.32 | 14.06 | No | N/A | No growth | N/A |
| 4 | 32.43 | 28.98 | No | N/A | No growth | N/A |
| 4 | 35.46 | 4.42 | No | N/A | No growth | N/A |
| 6 | N/A | N/A | No | N/A | N/A | No lesions seen. |
| 6 | N/A | N/A | No | N/A | N/A | No lesions seen. |
| 6 | 33.01 | 16.39 | No | N/A | No growth | N/A |
| 6 | 33.96 | 7.59 | No | N/A | No growth | N/A |
| 6 | 34.15 | 7.38 | No | N/A | No growth | N/A |
| 10 | N/A | N/A | No | N/A | N/A | Mild subcutaneous oedema, scanty lymphocytes, no pathogens. |
| 10 | N/A | N/A | No | N/A | N/A | Mild focal subcutaneous haemorrhage and oedema. Scanty scattered lymphocytes. No pathogens |
| 10 | 28.28 | 171.27 | No | N/A | 5.10E+03 | N/A |
| 10 | 28.28 | 130.82 | No | N/A | No growth | N/A |
| 10 | 29.84 | 39.44 | No | N/A | 3.67E+03 | N/A |
| 12 | 19.66 | 1.62E+04 | Yes | Ulcer | No growth | Missing data |
| 13 | 13.85 | 3.55E+05 | Yes | Ulcer | 3.07E+06 | Mild dermal and moderate subcutaneous oedema, mononuclear cell infiltrates, multifocal skeletal muscle necrosis. Scattered AFB present in necrotic and oedematous areas. |
| 14 | 14.35 | 7.02E+05 | Yes | Ulcer | 9.03E+05 | Severe subcutaneous oedema, mild inflammatory cell infiltrates (macrophages, neutrophils, lymphocytes), multifocal connective tissue and skeletal muscle necrosis. Locally extensive area of full thickness necrosis and ulceration of the epidermis. Clusters and single AFB are present. |
| 14 | 15.02 | 3.22E+05 | Yes | Ulcer | 2.17E+06 | Moderate dermal and severe subcutaneous oedema with focal vasculopathy, mononuclear cell infiltrates and connective tissue and skeletal muscle necrosis and loss. Moderate numbers of AFB scattered within necrotic and oedematous tissues. Mild diffuse epidermal hyperplasia. |
| 14 | 13.84 | 8.34E+05 | Yes | Ulcer | 2.94E+06 | Multifocal epidermal erosion with serocellular crusting. Moderate dermal and subcutaneous oedema, mixed inflammatory infiltrates, multifocal necrotising vasculitis and multifocal necrosis of skeletal muscle and interstitial tissues. Multifocal clusters of AFB in necrotic tissues and blood vessels and scattered within the oedematous subcutaneous tissues. |
| 14 | 14.38 | 4.72E+05 | Yes | Ulcer | 1.51E+06 | Large artefact with loss of subcutaneous tissues. Severe subcutaneous oedema, locally extensive epidermal and dermal necrosis, 50% of the perimeter. Severe focal necrotising vasculitis. Multifocal subcutaneous and skeletal muscle necrosis. Macrophage, neutrophil and lymphocyte infiltrations. multifocal abundant clusters and scattered AFB. |
| 14 | 15.91 | 1.72E+05 | Yes | Ulcer | 1.31E+06 | Multifocal dermal and diffuse subcutaneous oedema, with mixed inflammatory cell infiltrates, severe vasculopathy associated with necrosis. Necrosis of scattered skeletal muscle fibres and clusters and scattered AFB. |
| 15 | 13.84 | 5.96E+05 | Yes | Ulcer | 3.54E+06 | Moderate to diffuse dermal and subcutaneous oedema with moderate mixed inflammatory cell infiltrations, macrophages, and lymphocytes, with scattered intra-cellular and interstitial AFB. Multifocal vasculopathy. Multifocal skeletal muscle necrosis. |
| 15 | 12.08 | 2.23E+06 | Yes | Ulcer | No growth | Moderate dermal and subcutaneous oedema and vasculopathy with multifocal skeletal muscle and interstitial necrosis. Infiltrations of mixed inflammatory cells, macrophages and lymphocytes. AFB within the necrotic foci and associated with vasculitis and subcutaneous tissues. |
| 15 | 15.44 | 2.05E+05 | Yes | Ulcer | No growth | Severe full-thickness epidermal necrosis and erosion. Severe diffuse subcutaneous oedema with inflammatory cell infiltrates with scattered acid-fast bacilli. Focal severe necrotising vasculitis with abundant AFB. Multifocal skeletal muscle and interstitial necrosis with clusters of AFB. |

**Table S3.** Clinical, microbiological and histopathological features: BALB/c mice, high dose.

| Time after challenge (weeks) | Mean IS2404 PCR Ct | GE /g of tissue | Lesion | Lesion type | CFU/g tissue | Histology report |
| --- | --- | --- | --- | --- | --- | --- |
| 4 | N/A | N/A | No | N/A | N/A | 1 focus of skeletal muscle bundle hypertrophy, incidental. |
| 4 | N/A | N/A | No | N/A | N/A | No lesions seen. |
| 4 | 31.66 | 44.75 | No | N/A | No growth | N/A |
| 4 | 31.62 | 41.93 | No | N/A | No growth | N/A |
| 4 | 31.34 | 57.68 | No | N/A | No growth | N/A |
| 6 | N/A | N/A | No | N/A | N/A | No lesions seen. |
| 6 | N/A | N/A | No | N/A | N/A | No lesions seen. |
| 6 | 30.28 | 81.66 | No | N/A | No growth | N/A |
| 6 | 32.15 | 25.57 | No | N/A | No growth | N/A |
| 6 | 30.81 | 52.67 | No | N/A | No growth | N/A |
| 10 | N/A | N/A | No | N/A | N/A | No lesions seen. |
| 10 | N/A | N/A | No | N/A | N/A | No lesions seen. |
| 10 | 30.27 | 30.38 | No | N/A | 2.00E+03 | N/A |
| 10 | 17.60 | 7.45E+04 | No | N/A | 1.00E+05 | N/A |
| 10 | 31.48 | 19.11 | No | N/A | 2.56E+02 | N/A |
| 13 | 13.63 | 8.12E+05 | Yes | Ulcer | 2.67E+06 | Marked subcutaneous oedema and multifocal epidermal ulceration and segmental epidermal erosion, hyperplasia and hyperkeratosis. Focal epidermal pustule. Multifocal necrosis of skeletal muscles and connective tissues. Moderate mononuclear cell infiltration. Multifocal AFB, mostly focused in necrotic areas but also identified in the bone marrow. |
| 14 | 13.23 | 1.12E+06 | Yes | Ulcer | 7.18E+05 | Severe diffuse dermal and subcutaneous oedema, multifocal subcutaneous and muscle necrosis, infiltrations of macrophages, lymphocytes, neutrophils. Multifocal AFB, including in a follicle. |
| 14 | 13.66 | 7.95E+05 | Yes | Ulcer | 2.89E+06 | Moderate to severe diffuse dermal and subcutaneous oedema, necrosis of skeletal muscles and subcutaneous connective tissues, mononuclear cell infiltrates and AFB present. |
| 14 | 13.19 | 8.79E+05 | Yes | Ulcer | 1.56E+06 | Moderate to severe segmental dermal and subcutaneous oedema with multifocal necrosis and necrosis of the skeletal muscle. Focal epidermal ulceration. Mononuclear cell infiltrates. Diffuse epidermal hyperplasia and hyperkeratosis. Multifocal aggregates of AFB and scattered single bacilli within subcutaneous oedema tissue. |
| 14 | 13.68 | 7.87E+05 | Yes | Ulcer | 1.22E+06 | Severe diffuse dermal and subcutaneous oedema, mononuclear cell infiltrates, epidermal hyperplasia and hyperkeratosis and necrosis of skeletal muscle and connective tissues. Numerous AFB in areas of necrosis and scattered in the subcutaneous tissue. |
| 15 | 12.38 | 1.72E+06 | Yes | Ulcer | 2.53E+06 | Marked epidermal ulceration-transmural necrosis with pustule formation. Moderate segmental epidermal hyperplasia and hyperkeratosis. Severe diffuse dermal and subcutaneous oedema with mixed inflammatory infiltrates of macrophages and lymphocytes. Multifocal mild to severe vasculopathy with focal necrotising leukocytoclastic vasculitis, associated with myriad of AFB. Multifocal necrosis with cellular debris and the presence of AFB. |
| 16 | 13.21 | 1.39E+06 | Yes | Ulcer | 7.67E+05 | Segmental epidermal hyperplasia and hyperkeratosis. Mild dermal and moderate to severe subcutaneous oedema, multifocal necrotising vasculitis with tissue necrosis and mixed inflammatory infiltrates. Multifocal skeletal muscle fibre necrosis and oedema. AFB are present in clusters in necrotic tissue, within affected blood vessels perivascular tissues, vascular wall and lumen, and scattered in the subcutaneous tissue. |
| 17 | 15.92 | 1.19E+05 | Yes | Ulcer | 1.07E+06 | Severe diffuse dermal and subcutaneous oedema, mild mixed inflammatory cell infiltrate, multifocal vasculopathy, multifocal necrosis of the subcutaneous connective tissue and skeletal muscles fibres. |
| 17 | 15.24 | 1.80E+05 | Yes | Ulcer | 1.45E+06 | Severe segmental, (50%) dermal and subcutaneous oedema with multifocal tissue necrosis and inflammatory cell infiltration. Focal large cluster of AFB in the necrotic tissue. |
| 17 | 11.78 | 1.44E+06 | Yes | Ulcer | 8.26E+06 | Severe locally extensive full-thickness epidermal necrosis with serocellular crusting and multifocal pustules. Severe subcutaneous oedema with moderate mixed neutrophil, lymphocyte and macrophage infiltrations and vasculopathy. Large foci of skeletal muscle and interstitial necrosis. Numerous clusters of AFB in association with necrotic tissues and within inflammatory cells in the subcutaneous oedema. |

**Table S4.** Clinical, microbiological and histopathological features: BALB/c mice, low dose.

| Time after challenge (weeks) | Mean IS2404 PCR Ct | GE /g of tissue | Lesion | Lesion type | CFU/g tissue | Histology report |
| --- | --- | --- | --- | --- | --- | --- |
| 4 | N/A | N/A | No | N/A | N/A | Incidental tendon degeneration, tendinosis. No pathogens. |
| 4 | N/A | N/A | No | N/A | N/A | Incidental tendon degeneration, tendinosis. No pathogens. |
| 4 | 37.42 | 1.32 | No | N/A | No growth | N/A |
| 4 | 31.78 | 39.25 | No | N/A | No growth | N/A |
| 4 | 33.35 | 17.15 | No | N/A | No growth | N/A |
| 6 | N/A | N/A | No | N/A | N/A | Mild focal subcutaneous mononuclear cell infiltration into adipose tissue. No other pathology. No pathogens. |
| 6 | N/A | N/A | No | N/A | N/A | No lesions seen. |
| 6 | 33.51 | 10.38 | No | N/A | No growth | N/A |
| 6 | 33.92 | 8.13 | No | N/A | No growth | N/A |
| 6 | 33.56 | 12.22 | No | N/A | No growth | N/A |
| 10 | N/A | N/A | No | N/A | N/A | No lesions seen. |
| 10 | N/A | N/A | No | N/A | N/A | There is loss of fascicle structures, not sure what the pathogenesis of this change is. No pathogens. Several artefacts in these tendons/fascicles. |
| 10 | 29.38 | 67.46 | No | N/A | 4.36E+03 | N/A |
| 10 | 29.98 | 54.38 | No | N/A | No growth | N/A |
| 10 | 21.76 | 8.63E+03 | No | N/A | 5.88E+02 | N/A |
| 14 | 15.34 | 2.42E+05 | Yes | Ulcer | 2.38E+06 | Severe diffuse dermal and subcutaneous oedema with mild macrophage, lymphocyte, neutrophil infiltrations. Focal vasculopathy. Multifocal necrosis of skeletal muscle and connective tissue. Numerous foci of AFB. |
| 14 | 15.46 | 2.69E+05 | Yes | Ulcer | 2.78E+05 | Moderate diffuse subcutaneous oedema with mild inflammatory infiltrates, multifocal muscle necrosis and oedema. Multifocal AFB. Multifocal epidermal hyperplasia and hyperkeratosis. |
| 14 | 14.36 | 3.48E+05 | Yes | Ulcer | 1.76E+06 | Moderate diffuse subcutaneous oedema, scant inflammatory cell infiltration, vasculopathy, necrosis of connective tissue and skeletal muscle. Multifocal epidermal hyperplasia and hyperkeratosis. Multifocal AFB. |
| 16 | 13.93 | 1.02E+06 | Yes | Ulcer | 7.06E+06 | Severe locally extensive full-thickness necrosis and inflammation of the epidermis, dermis and subcutaneous tissues. Diffuse dermal and subcutaneous oedema with vasculopathy and mixed inflammatory infiltrates. Multifocal deep subcutaneous and skeletal muscle necrosis and inflammation. Abundant clusters and scattered AFB in the subcutaneous and affected muscle. |
| 17 | 15.34 | 3.38E+05 | Yes | Ulcer | 4.75E+05 | Severe diffuse dermal and subcutaneous oedema with mixed inflammatory cell infiltrates, and necrosis. Severe multifocal necrotising vasculitis with AFB within the perivascular tissues, vascular wall and lumen. Skeletal muscle necrosis and oedema. Numerous clusters and scattered AFB in affected tissues. |
| 17 | 16.17 | 2.50E+05 | Yes | Ulcer | 1.26E+04 | Mild segmental epidermal hyperplasia, hyperkeratosis and moderate loss of epidermal appendages Severe diffuse dermal and subcutaneous oedema, inflammatory cell infiltrate and necrosis. Multifocal necrotising vasculitis. Multifocal skeletal muscle necrosis. Multifocal clusters and scattered AFB bacteria are present. |
| 17 | 15.68 | 2.55E+05 | Yes | Ulcer | 2.36E+06 | Moderate segmental epidermal hyperplasia and hyperkeratosis. Severe diffuse dermal and subcutaneous oedema, inflammatory cell infiltrate and necrosis. Severe multifocal necrotising vasculitis with acid-fast bacilli within the perivascular tissues, vascular wall, and lumen. Multifocal skeletal muscle necrosis. Multifocal clusters and scattered single AFB are present. |
| 17 | 15.94 | 1.18E+05 | Yes | Ulcer | 3.05E+06 | Mild to moderate subcutaneous oedema and multifocal tissue necrosis. Moderate chronic active inflammation. Scant clusters and scattered AFB. |
| 19 | 16.32 | 1.61E+05 | Yes | Ulcer | 3.04E+06 | Moderate diffuse dermal and subcutaneous oedema, multifocal mild vasculopathy, multifocal necrosis of skeletal muscle and connective tissues, mild mixed macrophage and lymphocyte infiltrations. Focal cluster of acid-fast bacilli close to necrotic tissue, scanty AFB in oedematous subcutaneous tissue. |
| 20 | 15.95 | 8.36E+04 | Yes | Ulcer | 1.60E+06 | Moderate dermal and subcutaneous oedema with multifocal necrosis of skeletal muscle fibres and interstitium with mixed inflammatory infiltrates of macrophages, lymphocytes and lesser numbers of neutrophils. There is segmental epidermal hyperplasia and hyperkeratosis. AFB are present in clusters with necrotic foci and scattered in low numbers within the oedematous tissues. |

**Table S5.** Clinical and biometric features of C57BL/6 mice with lesions, and representative controls.

| High dose |  |  |  | Low dose |  |  |  |
| --- | --- | --- | --- | --- | --- | --- | --- |
| Mouse ID | Incubation (days) | Lesion onset to ulcer (days) | Change in weight (g) | Mouse ID | Incubation (days) | Lesion onset to ulcer (days) | Change in weight (g) |
| 8.1 | 64 | 13 | 1.0 | 19.1 | 77 | 18 | 1.5 |
| 8.2 | 60 | N/A | N/A | 19.2 | 81 | 24 | 1.7 |
| 8.3 | 60 | 6 | -1.2 | 19.3 | 77 | 25 | 5.6 |
| 8.4 | 67 | 14 | 0.1 | 19.4 | 72 | 23 | -0.2 |
| 8.5 | 62 | 15 | 0.4 | 19.5 | 86 | 14 | 0.7 |
| 9.1 | 63 | N/A | N/A | 20.1 | 72 | 13 | 1.3 |
| 9.2 | 72 | 16 | 1.1 | 20.2 | 76 | 15 | 0.8 |
| 9.3 | 72 | 19 | -0.2 | 20.3 | 71 | 25 | 1.8 |
| 9.4 | 65 | 18 | 1.5 | 20.4 | 74 | 22 | 0.7 |
| 9.5 | 63 | N/A | N/A | 20.5 | 85 | 17 | 2.3 |
| 10.1 | 64 | 13 | -0.3 | Mean: | 77 | 20 | 1.6* |
| 10.2 | 62 | 15 | -0.9 |  |  |  |  |
| 10.3 | 53 | 13 | 3.9 |  |  |  |  |
| 10.4 | 67 | 10 | 0.8 |  |  |  |  |
| 10.5 | 67 | 15 | 0.3 |  |  |  |  |
| Mean: | 64 | 14 | 0.5* |  |  |  |  |

| Negative controls |  |  |  | Negative controls |  |  |  |
| --- | --- | --- | --- | --- | --- | --- | --- |
| Mouse ID | Incubation (days) | Lesion onset to ulcer (days) | Change in weight (g) | Mouse ID | Incubation (days) | Lesion onset to ulcer (days) | Change in weight (g) |
| 9.1 | N/A | N/A | 0.4 | 9.2 | N/A | N/A | 1.0 |
| 9.2 | N/A | N/A | 0.1 | 9.3 | N/A | N/A | 1.3 |
| 9.3 | N/A | N/A | -0.5 | 9.4 | N/A | N/A | 1.1 |
| 9.4 | N/A | N/A | 0.0 | 9.5 | N/A | N/A | 1.2 |
| 9.5 | N/A | N/A | -0.2 | 10.1 | N/A | N/A | 0.8 |
| 10.1 | N/A | N/A | 0.4 | 10.2 | N/A | N/A | 0.7 |
| 10.2 | N/A | N/A | 0.3 | 10.3 | N/A | N/A | 3.5 |
| 10.3 | N/A | N/A | 0.3 | 10.4 | N/A | N/A | 0.1 |
| 10.4 | N/A | N/A | 0.8 | 10.5 | N/A | N/A | 0.9 |
| 10.5 | N/A | N/A | 0.0 | Mean: | N/A | N/A | 1.2* |
| Mean: | N/A | N/A | 0.2* |  |  |  |  |

\*Student's t test, p = 0.382
\*Student's t test, p = 0.442

**Table S6.** Clinical and biometric features of BALB/c mice with lesions, and representative controls.

| High dose |  |  |  | Low dose |  |  |  |
| --- | --- | --- | --- | --- | --- | --- | --- |
| Mouse ID | Incubation (d) | Lesion onset to ulcer (days) | Change in weight (g) | Mouse ID | Incubation (d) | Lesion onset to ulcer (days) | Change in weight (g) |
| 4.1 | 85 | 24 | 1.0 | 13.1 | 67 | 31 | 0.6 |
| 4.2 | 91 | 27 | 1.0 | 13.2 | 103 | 36 | 2.5 |
| 4.3 | 82 | 20 | -0.4 | 13.3 | 93 | 25 | 0.6 |
| 4.4 | 83 | 34 | 0.7 | 13.4 | 72 | 26 | 1.1 |
| 4.5 | 89 | 32 | 3.3 | 13.5 | 86 | 32 | -0.1 |
| 5.1 | 72 | 25 | 0.7 | 14.1 | 81 | 37 | 0.2 |
| 5.2 | 73 | 19 | 0.9 | 14.2 | 105 | 29 | 1.5 |
| 5.3 | 72 | 23 | 1.6 | 14.3 | 73 | 25 | 0.9 |
| 5.4 | 72 | 23 | 1.3 | 14.4 | 79 | 38 | 1.8 |
| 5.5 | 72 | 26 | 0.8 | 14.5 | 79 | 35 | 0.5 |
| Mean: | 79 | 25 | 1.1* | Mean: | 84 | 32 | 1.0* |
| Negative controls |  |  |  | Negative controls |  |  |  |
| 4.1 | N/A | N/A | 1.9 | 5.1 | N/A | N/A | 0.3 |
| 4.2 | N/A | N/A | 0.6 | 5.2 | N/A | N/A | 0.5 |
| 4.3 | N/A | N/A | 1.3 | 5.3 | N/A | N/A | 1.7 |
| 4.4 | N/A | N/A | 1.0 | 5.4 | N/A | N/A | 1.3 |
| 4.5 | N/A | N/A | 1.5 | 5.5 | N/A | N/A | 1.6 |
| 5.1 | N/A | N/A | 1.4 | Mean: | N/A | N/A | 1.1* |
| 5.2 | N/A | N/A | 0.2 | *Student's t test, p = 0.773 |  |  |  |
| 5.3 | N/A | N/A | 1.0 |  |  |  |  |
| 5.4 | N/A | N/A | 0.8 |  |  |  |  |
| 5.5 | N/A | N/A | 1.5 |  |  |  |  |
| Mean: | N/A | N/A | 1.1* |  |  |  |  |
| *Student's t test, p = 0.930 |  |  |  |  |  |  |  |

**Table S7.** Statistically significant differences between *M. ulcerans* dosing groups (high dose, low dose, and control) and various immune features over time (unadjusted univariate analysis).

| Parameter | Timepoint (weeks) | Location | Higher value | Lower value | p value |
| --- | --- | --- | --- | --- | --- |
| CD8<sup>+</sup> cells | 4 | Spleen | Low dose | Control | 0.016 |
|  | 4 | Spleen | Low dose | High dose | 0.016 |
|  | 10 | DLN | Low dose | Control | 0.036 |
|  | > 10 | DLN | Low dose | Control | 0.021 |
| CD4<sup>+</sup> cells | 4 | Spleen | Low dose | Control | 0.016 |
|  | 4 | Spleen | Low dose | High dose | 0.016 |
|  | > 10 | DLN | Low dose | Control | 0.017 |
| T<sub>reg</sub> | 4 | Spleen | Low dose | Control | 0.016 |
|  | 4 | Spleen | Low dose | High dose | 0.016 |
|  | 10 | Spleen | Control | Low dose | 0.036 |
|  | > 10 | DLN | Low dose | Control | 0.008 |
| B cells | 4 | Spleen | Low dose | Control | 0.016 |
|  | 4 | Spleen | Low dose | High dose | 0.016 |
|  | 10 | DLN | Low dose | Control | 0.036 |
| Plasma cells | 4 | Spleen | Low dose | Control | 0.016 |
|  | 4 | Spleen | Low dose | High dose | 0.032 |
|  | 6 | DLN | Low dose | Control | 0.008 |
|  | >10 | DLN | Low dose | Control | 0.004 |
| γδ T cells | 4 | Spleen | Low dose | Control | 0.016 |
|  | 4 | Spleen | Low dose | High dose | 0.016 |
|  | 6 | DLN | High dose | Control | 0.032 |
|  | > 10 | DLN | Low dose | Control | 0.002 |
|  | > 10 | DLN | Low dose | High dose | 0.029 |
| NK cells | 4 | Spleen | Low dose | Control | 0.016 |
|  | 6 | DLN | Low dose | Control | 0.008 |
|  | 6 | DLN | High dose | Control | 0.016 |
|  | > 10 | DLN | Low dose | Control | 0.017 |
|  | > 10 | DLN | Low dose | High dose | 0.019 |
| Macrophages | 4 | Spleen | Low dose | Control | 0.016 |
|  | 4 | Spleen | Low dose | High dose | 0.016 |
|  | 6 | DLN | Low dose | Control | 0.025 |
|  | 6 | DLN | High dose | Control | 0.025 |
|  | 10 | DLN | Low dose | Control | 0.036 |
|  | > 10 | DLN | High dose | Control | 0.036 |
|  | > 10 | DLN | Low dose | Control | <0.001 |
|  | > 10 | DLN | Low dose | High dose | <0.001 |
| Neutrophils | 4 | Spleen | Low dose | High dose | 0.016 |
|  | 10 | DLN | Low dose | Control | 0.036 |
|  | > 10 | Spleen | Low dose | Control | <0.001 |
| Dendritic cells | 4 | Spleen | Low dose | Control | 0.016 |
|  | 4 | Spleen | Low dose | High dose | 0.032 |
|  | 6 | DLN | Low dose | Control | 0.032 |
|  | > 10 | DLN | Low dose | Control | 0.003 |
|  | > 10 | DLN | Low dose | High dose | 0.012 |
| IL-10 | 4 | Serum | Control | High dose | 0.012 |
| CXCL10 | 6 | Serum | Low dose | Control | 0.012 |
| CCL2 (MIP1) | 6 | Serum | Low dose | Control | 0.034 |
|  | > 10 | Serum | Low dose | Control | 0.001 |
|  | > 10 | Serum | High dose | Control | 0.006 |
| CXCL9 | > 10 | Serum | High dose | Control | 0.045 |
| VEGF | 4 | Serum | High dose | Control | 0.045 |
|  | 10 | Serum | Low dose | Control | 0.049 |
| IL-4 | 6 | Serum | Low dose | Control | 0.016 |
| IL-6 | > 10 | Serum | High dose | Control | 0.002 |
|  | > 10 | Serum | High dose | Low dose | 0.005 |
| IFN-α | > 10 | Serum | High dose | Control | 0.019 |

BALB/c
| Parameter | Timepoint (weeks) | Location | Higher value | Lower value | p value |
| --- | --- | --- | --- | --- | --- |
| CD8<sup>+</sup> cells | 10 | Spleen | Control | Low dose | 0.008 |
|  | 10 | Spleen | Control | High dose | 0.032 |
| CD4<sup>+</sup> cells | 10 | Spleen | Control | Low dose | 0.016 |
|  | 10 | Spleen | Control | High dose | 0.008 |
| T<sub>reg</sub> | 10 | Spleen | Control | Low dose | 0.016 |
|  | 10 | Spleen | Control | High dose | 0.008 |
| B cells | 6 | DLN | Control | Low dose | 0.032 |
|  | 10 | Spleen | Control | Low dose | 0.008 |
|  | 10 | Spleen | Control | High dose | 0.016 |
| Plasma cells | 4 | DLN | Control | Low dose | 0.032 |
|  | 6 | DLN | Control | Low dose | 0.032 |
|  | 10 | Spleen | Control | Low dose | 0.032 |
| γδ T cells | 6 | DLN | Control | Low dose | 0.032 |
|  | 10 | Spleen | Control | Low dose | 0.016 |
|  | 10 | Spleen | Control | High dose | 0.032 |
| NK cells | 4 | DLN | Control | High dose | 0.032 |
|  | 10 | Spleen | Control | Low dose | 0.016 |
|  | 10 | Spleen | Control | High dose | 0.008 |
|  | 10 | DLN | Control | Low dose | 0.008 |
|  | 10 | DLN | Control | High dose | 0.008 |
| Macrophages | 4 | DLN | Control | High dose | 0.042 |
|  | 10 | Spleen | Control | Low dose | 0.008 |
|  | 10 | Spleen | Control | High dose | 0.008 |
| Neutrophils | 10 | Spleen | Control | Low dose | 0.016 |
| Dendritic cells | 10 | Spleen | Control | Low dose | 0.008 |
|  | 10 | Spleen | Control | High dose | 0.008 |
| IFN-α | 4 | Serum | Control | Low dose | 0.016 |
| IFN-γ | 6 | Serum | High dose | Low dose | 0.045 |
| CXCL9 | 4 | Serum | High dose | Low dose | 0.030 |
| GM-CSF | 6 | Serum | Control | Low dose | 0.025 |
| IL-10 | 10 | Serum | High dose | Control | 0.025 |
|  | 10 | Serum | High dose | Low dose | 0.044 |

**Table S8.** p values reaching statistical significance following correction for multiple analyses across the entire univariate dataset. C57BL/6 mice only are presented.

| Parameter | Timepoint (weeks) | Location | Higher value | Lower value | p value | Adj. p value |
| --- | --- | --- | --- | --- | --- | --- |
| CD44<sup>+</sup> CD62L<sup>lo</sup> CD4<sup>+</sup> | > 10 | DLN | Low dose | Control | 0.002 | 0.038 |
| CD38<sup>+</sup> CD8<sup>+</sup> | > 10 | DLN | Low dose | Control | 0.003 | 0.038 |
| CD38<sup>+</sup> CD8<sup>+</sup> | > 10 | DLN | Low dose | High dose | 0.002 | 0.048 |

**Table S9.**
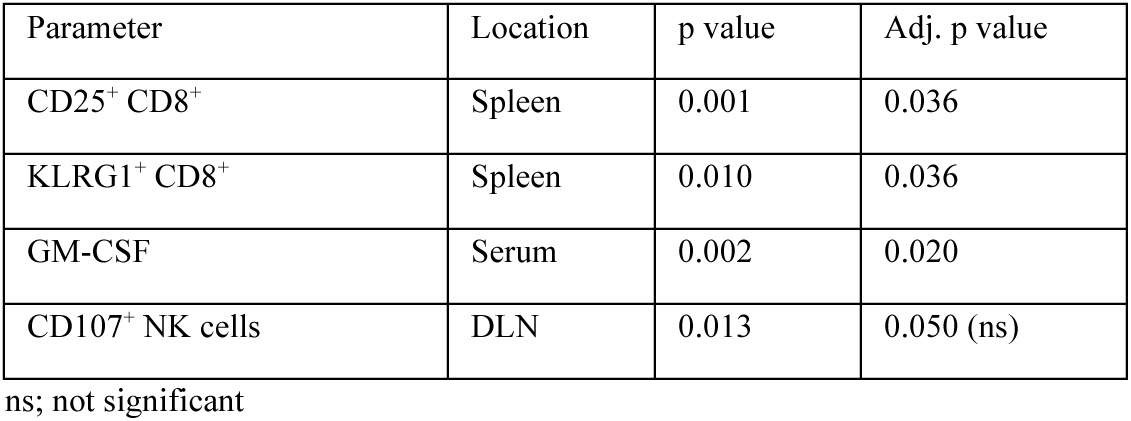
p values reaching statistical significance following correction for multiple analyses across the dataset presented in Fig 5E. C57BL/6 mice only are presented.

